# c-di-AMP determines the hierarchical organization of bacterial RCK proteins

**DOI:** 10.1101/2023.04.24.538052

**Authors:** Rita Rocha, João M. P. Jorge, Celso M. Texeira-Duarte, Inês R. Figueiredo-Costa, Christina Herzberg, Jörg Stülke, João H. Morais-Cabral

## Abstract

RCK domains or proteins are important regulatory components of cation and K^+^ channels and transporters both in eukaryotic and prokaryotic organisms. A lasting unanswered question about bacterial RCK proteins relates to the physiological role of the multiple, sometimes closely related, RCK genes encoded in the genome. We explored this question with the Ktr channels of *Bacillus subtilis* that include two genes encoding RCK proteins (KtrA and KtrC) and two genes encoding their membrane protein partners (KtrB and KtrD). Using a combination of *in vivo* characterization and *in vitro* functional analysis, we determined that the two RCK proteins are neither physiologically redundant or functionally equivalent. Instead, KtrC is the physiologically dominant RCK protein due to its ability to mediate K^+^ transport inactivation by c-di-AMP, a bacterial second messenger that is the master regulator of the K^+^ machinery in many species, while KtrA assembled channels are very insensitive to the dinucleotide. Moreover, KtrC and KtrA can form heteromeric assemblies that can control the Ktr channel activity and are sensitive to c-di-AMP inhibition. In parallel, we showed that conditions with a large number of Ktr channels assembled with KtrA, or with RCK proteins that do not mediate c-di-AMP inhibition, are toxic to the cell. Altogether, we have demonstrated that c-di-AMP regulation of Ktr channels goes beyond affecting transcription and functional activity, it also determines the hierarchical organization of bacterial RCK proteins.

## Introduction

Like all cells, bacteria accumulate K^+^ (1). However, unlike mammalian cells, bacteria are able to vary their intracellular K^+^ levels, using it to regulate intracellular pH (2-4), turgor (2, 5, 6) and membrane potential (7). For example, in the early phase of adaptation to hyperosmotic shock, intracellular K^+^ in *B. subtilis* rises rapidly from ∼300 mM to 600-700 mM, before dropping back to lower levels as other adaptation mechanisms come into play (8). The K^+^ importers and exporters (or the K^+^ machinery) that underlie these variations are known and belong to conserved families (7, 9, 10). One of these is the Ktr cation channel family (11-13). These bacterial channels are widespread and composed by a membrane protein and a regulatory protein, a Regulator of Conductance of K^+^ (RCK) domain. RCK domains are present in eukaryotic and prokaryotic organisms where they modulate the activity of cation channels and transporters (14, 15). In bacteria and archaea RCK domains constitute more than half of the regulatory domains of identified prokaryotic K^+^ channels (16) and they exist both as domains of a larger membrane protein polypeptide or as separate proteins, which we will refer to as RCK proteins. Importantly, the second messenger c-di-AMP, which has been proposed to regulate intracellular K^+^ in many bacterial species, is thought to regulate Ktr channel function by binding to RCK proteins (17).

The genomic and functional organization of bacterial RCK proteins and their Ktr membrane protein partners is diverse and relatively complex. For example, *Staphylococcus aureus* encodes a single RCK protein that is thought to assemble with the Ktr membrane proteins from two different genes, *Salinicoccus halodurans* contains genes for two RCK protein and a single Ktr membrane protein and *B. subtilis* encodes two RCK proteins (KtrA and KtrC) and two Ktr membrane proteins (KtrB and KtrD). The complexity in *B. subtilis* is particularly evident, with the gene of KtrA forming an operon with the gene of KtrB while the gene for KtrC is in an operon with *kinC* and *ykqA*, coding for proteins unrelated to Ktr channels, a sensor kinase of the Spo0A phosphorelay and a putative gamma-glutamylcyclotransferase, respectively. In contrast, KtrD is encoded by a stand-alone gene. The two RCK proteins (KtrA and KtrC) are closely related, sharing 55% amino acid sequence identity. They both display the canonical tertiary organization of RCK proteins, with an N- and a C-terminal subdomain The two Ktr membrane proteins, KtrB and KtrD, are also very similar, with 36% amino acid sequence identity. Assembly of the RCK proteins with the Ktr membrane protein is essential for channel activity and crystal and cryo-EM structures of KtrAB complexes from *B. subtilis* and *Vibrio alginolyticus* have revealed a KtrB channel dimer associated with a cytosolic KtrA octameric ring (13, 18, 19). Partly due to pairing of the *ktrA* and *ktrB* genes in the same *B. subtilis* operon, it has been proposed that in the cell, these two proteins assemble as KtrAB while KtrC assembles with KtrD, forming KtrCD (11). However, we have recently demonstrated that the two RCK proteins are promiscuous and are able to activate both KtrB and KtrD (12, 20). Finally, while the N-terminal subdomains of KtrA and KtrC include a binding site for ATP (activator) or ADP (inactivator) (13, 19, 21), c-di-AMP binds to the C-terminal subdomains where it is thought to act as an inactivator (17, 22). The constants of dissociation of c-di-AMP to the isolated KtrA and KtrC proteins were determined as ∼3 μM and ∼30 nM, respectively (20). Additionally, transcription of the *ktrAktrB* operon is down-regulated by c-di-AMP that binds to a riboswitch positioned just after the operon’s promoter (23, 24).

Despite the extensive characterization of the molecular properties of Ktr channels, there is little or no understanding about the physiological function of their relatively complex organization, and in particular for the need of multiple but very similar regulatory RCK proteins in a bacterium. Moreover, the role of K^+^ machinery master regulator c-di-AMP in the function of individual RCK proteins has not been fully established and the impact of the dinucleotide in defining their physiological role is yet to be explored.

To gain insights into these questions we have analyzed the functional and physiological properties and relationships of the RCK proteins and their membrane protein partners in the model Gram-positive organism *B. subtilis*. Our results demonstrate that the functional organization of RCK proteins in *B. subtilis* is more complex than previously thought and that c-di-AMP has a crucial role in determining the regulatory function of individual RCK proteins as well as in defining their physiological hierarchy, representing a novel understanding of the role of RCK domains in bacteria.

## Material and methods

### Protein Expression and Purification

Tag-less wild-type KtrA cloned in a modified pET-24d vector, tag-less wild-type KtrC cloned in pET24a vector and respective mutants were over-expressed in the *Escherichia coli* BL21 (DE3) strain. Cells transformed with an expression vector were grown at 37 ºC with agitation in LB medium supplemented with 50 mg/L of kanamycin to an OD_600_ = 0.6-0.8. Protein expression was induced for 14-16 h at 20 ºC with 0.5 mM IPTG, after immersion of cultures in ice for 30 min and addition of 1 or 2 % (v/v) ethanol, for KtrC or KtrA, respectively. Cell lysis was done in 50 mM Tris-HCl pH 7.5, 50 mM KCl, 5 mM DTT supplemented with protease inhibitors (1 mM PMSF, 1 µg/mL leupeptin, 1 µg/mL pepstatin A) using a cooled cell cracker Emulsiflex-C5 (Avestin). Lysate was cleared by centrifugation and the supernatant loaded into an anion exchange column Hi-Trap Sepharose Q-HP (GE Healthcare). KtrA and KtrC were eluted with a 50-1000 mM KCl gradient. Protein-containing fractions were pooled and incubated with an ATP-agarose resin (Immobilized γ-Aminophenyl-ATP C10-spacer from Jena Bioscience) overnight at 4 °C with gentle agitation. The resin was then washed thoroughly with 50 mM Tris-HCl pH 8.0, 150 mM KCl, 1 mM TCEP for KtrA or 50 mM Tris-HCl pH 8.0, 150 mM KCl, 5 mM DTT for KtrC and proteins were eluted in wash buffer supplemented with 5 mM ATP or ADP (sodium salts). Proteins were concentrated and further purified by size-exclusion chromatography in a Superdex-S200 (GE Healthcare) column using 50 mM Tris-HCl pH 8.0, 150 mM KCl, 5 mM DTT. KtrA and KtrC were then supplemented with 1 mM ATP or ADP. For addition to functional assays, the size-exclusion chromatography step was done in 50 mM Tris-HCl pH 8.0, 150 mM KCl, 5 mM DTT, 1 mM MgCl_2_ and proteins were supplemented with 0.1 mM ATP or ADP. Protein concentration was determined using a colorimetric assay (Bio-Rad).

N-terminal Strep-tagged KtrA cloned in a modified pET-24d vector was over-expressed in the *Escherichia coli* BL21 (DE3) strain. Cells transformed with the expression vector were grown at 37 ºC with agitation in LB medium supplemented with 50 mg/L of kanamycin to an OD_600_ = 0.6-0.8. Protein expression was induced for 14-16 h at 20 ºC with 0.5 mM IPTG, after immersion of the cultures in ice for 30 min and addition of 1 % (v/v) ethanol. Cell lysis was done in 50 mM Tris-HCl pH 8.0, 150 mM KCl, 5 mM DTT supplemented with protease inhibitors (1 mM PMSF, 1 µg/mL leupeptin, 1 µg/mL pepstatin A) using a cooled cell cracker Emulsiflex-C5 (Avestin). Lysate was cleared by centrifugation and the supernatant was incubated with a Strep-Tactin Sepharose resin (IBA Lifesciences). Resin was thoroughly washed with lysis buffer and 50 mM Tris-HCl pH 8.0, 300 mM KCl, 5 mM DTT and the protein was eluted with lysis buffer supplemented with 10 mM desthiobiotin. Eluted protein was incubated with an ATP-agarose resin (Immobilized γ-Aminophenyl-ATP C10-spacer from Jena Bioscience) overnight at 4 °C with gentle agitation. The resin was washed thoroughly with 50 mM Tris-HCl pH 8.0, 150 mM KCl, 5 mM DTT and protein was eluted in wash buffer supplemented with 5 mM ATP (sodium salt). For addition to functional assays, protein was concentrated and further purified by size-exclusion chromatography in a Superdex-S200 (GE Healthcare) column equilibrated with 50 mM Tris-HCl pH 8.0, 150 mM KCl, 5 mM DTT, 1 mM MgCl_2_ and supplemented with 0.1 mM ATP. Protein concentration was determined using with a colorimetric assay (Bio-Rad).

C-terminal His-tagged KtrC cloned in pET24d was over-expressed in the *Escherichia coli* BL21 (DE3) strain. Cells transformed with the expression vector were grown at 37 ºC with agitation in LB medium supplemented with 50 mg/L of kanamycin to an OD_600_ = 0.6-0.8. Protein expression was induced for 14-16 h at 20 ºC with 0.5 mM IPTG, after immersion of the cultures in ice for 30 min and addition of 1 % (v/v) ethanol. Cell lysis was done in 50 mM Tris-HCl pH 8.0, 150 mM KCl, 5 mM DTT supplemented with protease inhibitors (1 mM PMSF, 1 µg/mL leupeptin, 1 µg/mL pepstatin A) using a cooled cell cracker Emulsiflex-C5 (Avestin). Lysate was cleared by centrifugation and the supernatant was incubated with Ni^2+^-beads, washed first with lysis buffer and then lysis buffer with 10 mM and 20 mM Imidazole. Protein was eluted with lysis buffer with 150 mM Imidazole and 1 mM ATP. Protein was further purified by size-exclusion chromatography in a Superdex-S200 (GE Healthcare) column equilibrated with 50 mM Tris-HCl pH 8.0, 150 mM KCl, 5 mM DTT, 1 mM MgCl_2_ and supplemented with 0.1 mM ATP. Protein concentration was determined using a colorimetric assay (Bio-Rad).

N-terminal Strep-tagged KtrB was over-expressed in the *Escherichia coli* BL21 (DE3) strain. Cells transformed with an expression vector were grown at 37 ºC with agitation in LB medium supplemented with 50 mg/L of kanamycin to an OD_600_ = 0.9-1.0. Protein expression was induced at 37 °C for 2.5 h with the addition of 0.5 mM IPTG and 1 mM BaCl_2_. Cells were lysed in 50 mM Tris-HCl pH 8.0, 120 mM NaCl, 30 mM KCl supplemented with protease inhibitors (1 mM PMSF, 1 µg/mL leupeptin, 1 µg/mL pepstatin A) using a cooled cell cracker Emulsiflex-C5 (Avestin). KtrB was extracted with 40 mM n-dodecyl-β-D-maltoside (DDM) (Sol-grade from Anatrace) overnight at 4 °C with gentle agitation. Spin-cleared lysate was loaded into a Strep-Tactin Sepharose resin (IBA Lifesciences) and washed with 50 mM Tris-HCl pH 8.0, 120 mM NaCl, 30 mM KCl, 1 mM DDM, 5 mM DTT, 1 mM ATP or ADP.

N-terminal His-tagged KtrD cloned into pRSFDuet-1 (Novagen) was overexpressed in Escherichia coli BL21 (DE3) grown in LB media supplemented with kanamycin 50 µg/mL. KtrD expression was induced at late-exponential phase with 0.5 mM IPTG for 2 hours at 37°C, in the presence of 1 mM BaCl_2_. Bacterial cells were lysed in 50 mM Tris-HCl pH 8.0, 120 mM NaCl, 30 mM KCl supplemented with protease inhibitors and KtrD was extracted overnight at 4°C with 40 mM DDM (n-Dodecyl-β-D-Maltopyranoside). Cell lysate was cleared by centrifugation (34,957 x g for 45 min at 4°C) and 5 mM Imidazole was added to the cell lysate, which was loaded into Co^2+^ resin (Talon) column pre-equilibrated with 50 mM Tris-HCl pH 8.0, 120 mM NaCl, 30 mM KCl, 1 mM DDM supplemented with 5 mM Imidazole. Co^2+^ resin was washed equilibration buffer supplemented with 5 mM Imidazole and next with same buffer plus 20 mM Imidazole. KtrD was eluted in equilibration buffer supplemented with 150 mM Imidazole and immediately diluted three times in the same buffer with 15 mM DTT.

To assemble the KtrAB or KtrCB complexes, purified KtrA or KtrC (with ATP or ADP) was added to Streptactin resin with bound KtrB and incubated for 30 min at 4 ºC with gentle agitation. The resin was thoroughly washed with 50 mM Tris-HCl pH 8.0, 120 mM NaCl, 30 mM KCl, 1 mM DDM, 5 mM DTT, 1 mM ATP or ADP and complexes were eluted with same buffer supplemented with 5 mM desthiobiotin. The eluted protein was dialysed overnight at 4 ºC against Dialysis buffer 50 mM Tris-HCl pH 8.0, 120 mM NaCl, 30 mM KCl, 0.5 mM DDM, 5 mM DTT, in the presence of thrombin for KtrB tag cleavage. For proteoliposome reconstitution, dialysis was performed against Internal Buffer (10 mM HEPES, 7 mM N-methyl-D-glucamine (NMG) (pH 8.0), 150 mM KCl) supplemented with 0.5 mM DDM and 5 mM DTT. Protein was further purified by size-exclusion chromatography in a Superdex-S200 (GE Healthcare) column using Dialysis buffer or Internal Buffer supplemented with 0.5 mM DDM and 5 mM DTT. Eluted fractions corresponding to KtrAB or KtrCB were pooled, concentrated to 1 mg/mL and used for proteoliposome preparation when required. No nucleotide was added during dialysis and size-exclusion chromatography. Protein concentration was estimated by measuring absorbance at 280 nm on a NanoDrop ND-1000 Spectrophotometer (NanoDrop Technologies). KtrB alone was purified in the same manner as KtrAB and KtrCB complexes but without the KtrA or KtrC addition step.

For protein co-assembly tests in *E. coli*, gene pairs were cloned into the two multiple cloning sites of pRSFDuet adding a Strep tag to one of the proteins and a His tag (or no tag) to the other. Cultures of BL21(DE3) *E. coli* transformed with plasmid were grown: for RCK proteins, to OD_600_= 0.6-0.8 with protein expression induced for 14-16 h at 20 ºC with 0.5 mM IPTG (after immersion of the cultures in ice for 30 min and addition of 1 % (v/v) ethanol); for KtrB/KtrD, to OD_600_ = 0.9-1.0 with protein expression induced at 37 °C for 2.5 h with the addition of 0.5 mM IPTG and 1 mM BaCl_2_. Pull-down experiments followed purification procedures described for Strep-tagged KtrA, His-tagged KtrC, Strep-tagged KtrB and His-tagged KtrD. For co-assembly *in vitro*, proteins were purified separately as described above and mixed 1:1 at 0.5 mg/mL final concentration in the presence of 0.1 mM ATP, ADP or c-di-AMP. Pull-down at different time points was as above. Pulled-down fractions were analysed by SDS-PAGE or Western blot using rabbit antibodies against KtrB or mouse against His-tag (Qiagen).

For assembly of N-terminal Strep-tagged KtrA/KtrC hetero-rings for functional assays with liposomes, ATP-bound N-terminal Strep-tagged KtrA and KtrC were purified separately until the elution step of the ATP-agarose resin, described above. At this point, both proteins were supplemented with 5 mM MgCl_2_ and protein concentration was determined using a colorimetric assay (Bio-Rad). Proteins were then mixed at a 2.5:1 mass ratio of KtrC to N-terminal Strep-tagged KtrA (11 mg of KtrC and 4.5 mg of N-terminal Strep-tagged KtrA in 10 mL total volume of 50 mM Tris-HCl pH 8.0, 150 mM KCl, 5 mM DTT, 1 mM MgCl_2,_ 0.1 mM ATP and incubated for 48h at 4ºC with gentle agitation. The mixture was then incubated with a Strep-Tactin Sepharose resin (IBA Lifesciences). Resin was thoroughly washed with the same buffer without ATP and protein was eluted with buffer supplemented with 5 mM desthiobiotin and 0.1 mM ATP. Eluted protein was further purified by size-exclusion chromatography in a Superdex-S200 (GE Healthcare) column equilibrated with 50 mM Tris-HCl pH 8.0, 150 mM KCl, 5 mM DTT, 1 mM MgCl_2_, 0.1 mM ATP. Protein concentration was determined using a colorimetric assay (Bio-Rad).

### Preparation of proteoliposomes

Proteoliposome preparation followed previously described methods with some modifications (13, 19, 21). Briefly, *E. coli* polar lipids (Avanti) were resuspended at 10 mg/mL in Internal Buffer (150 mM KCl, 10 mM HEPES, 7 mM NMG (pH 8.0)) and solubilized by adding 40 mM DM (n-Decyl-β-D-Maltoside) (Sol-Grade from Anatrace). KtrB alone, KtrAB or KtrCB complexes were purified just before reconstitution, added to solubilized lipids at 1:100 (w:w) protein-to-lipid ratio and incubated for 30 min at room temperature. In control liposomes (empty), Internal Buffer was added to lipids instead of protein. Detergent was removed using adsorbing SM-2 Biobeads (Bio-Rad): the protein-lipid mix was incubated twice with fresh Biobeads at 10:1 (w:w) bead-to-detergent ratio at room temperature for 1 h and then incubated overnight at 4 ºC at a 20:1 (w:w) bead-to-detergent ratio. Proteoliposomes were then flash-frozen in liquid nitrogen and stored at -80 ºC.

### Pyranine encapsulation

Pyranine encapsulation was done as previously described (25). Proteoliposomes were thawed in a 37 ºC water bath for 15 min, briefly sonicated and mixed with an equal volume of Internal Buffer containing 500 µM pyranine, for a final pyranine concentration of 250 µM. The mixture was then subjected to three freeze/thaw cycles and extruded to homogeneity using a 400 nm polycarbonate membrane filter (Avanti Mini-Extruder). Non-encapsulated pyranine was removed through a PD-10 Desalting column (Cytiva), pre-incubated with Internal Buffer. Proteoliposomes were then concentrated by ultracentrifugation (30 min; 100.000 g; 12 ºC) in an Optima Max XP Tabletop Ultracentrifuge (Beckman Coulter), resuspended in Internal Buffer and kept at 12 ºC.

### ACMA-fluorescence-based K^+^ flux assay

The assay was performed as previously described (21, 26). Proteoliposomes were thawed in a 37 ºC water bath for 15 min, briefly sonicated and kept at room temperature. The proteoliposomes were then diluted 100-fold (10 µL sample in 1 mL final volume) in Flux Buffer (10 mM HEPES and 7 mM NMG (pH 8.0), 150 mM choline chloride, 0.1 mM ATP; 2 mM MgCl_2_) to establish the K^+^ gradient. In experiments involving RCK additions to KtrB proteoliposomes, 10 µg of N-Strep-tagged KtrA-ATP, KtrA-ATP or KtrC-ATP were incubated for 10 min with the KtrB proteoliposomes before establishing the K^+^ gradient. For control liposomes (empty) or KtrB alone, buffer solution used for the size-exclusion chromatography step of KtrB purification was added instead of protein. The pH-sensitive dye 9-amino-6-chloro-2-methoxyacridine (ACMA) was then added to the proteoliposomes to a final concentration of 550 nM. Fluorescence was monitored every 2 s (λ_Ex_=410 nm, λ_Em_=480 nm), using a 104F-QS 10 mm quartz cuvette (Hellma) with a small magnet in a Horiba FluoroMax-4 spectrofluorometer (Horiba Scientific). After measuring the initial baseline fluorescence during 100 s, the assay was initiated by adding 1 µM of the H^+^ ionophore carbonyl cyanide m-chlorophenyl hydrazine (CCCP) and the flux signal was monitored for 400 s. For each experimental condition, one additional replica was performed in which, after baseline fluorescence measurement and CCCP addition, 296 nM of the K^+^ ionophore valinomycin was quickly added and flux signal was monitored for an additional 100s. This was performed to avoid valinomycin contamination of the cuvette between sample measurements and was done every day for each proteoliposome preparation. Valinomycin incorporates into all liposomes and mediates K^+^ efflux, indicating the total flux of the liposome population. The maximum value of fluorescence quenching change obtained was used to normalize the data.

Fluorescence quenching curves were normalized individually as follows:

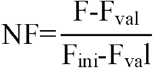

NF is the normalized fluorescence, F is the measured fluorescence in arbitrary units, F_ini_ corresponds to the last baseline point measured before CCCP addition and F_val_ corresponds to the lowest point measured after valinomycin addition.

### Pyranine-fluorescence-based K^+^ flux assay

The assay was performed as described by others with some modifications (25). 10 µL KtrB proteoliposomes were incubated with 4 µL (4 µg) RCK proteins and 2 µL of c-di-AMP or buffer (50 mM Tris-HCl pH 8.0, 150 mM KCl, 0.1 mM ATP, 1 mM MgCl_2_, 5 mM DTT) for 5 min. The mixture was then diluted 80-fold (16 µL sample in 1.3 mL final volume) in Flux Buffer (10 mM HEPES and 7 mM NMG (pH 8.0), 150 mM choline chloride, 0.1 mM ATP or ADP, 2 mM MgCl_2_) to establish the K^+^ gradient and incubated for 3 min inside the cuvette with agitation before starting the fluorescence measurements. Fluorescence was monitored every 2 s (λ_Ex_=460 nm, λ_Em_=510 nm), using a 104F-QS 10 mm quartz cuvette (Hellma) with a small magnet in a Horiba FluoroMax-4 spectrofluorometer (Horiba Scientific). After measuring the initial baseline fluorescence during 50 s, the assay was initiated by adding 4 µM of the H^+^ ionophore carbonyl cyanide m-chlorophenyl hydrazine (CCCP). Fluorescence was monitored for 450 s.

Fluorescence quenching curves were normalized individually as follows:

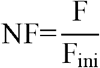

NF is the normalized fluorescence; F is the measured fluorescence in arbitrary units and F_ini_ corresponds to the average of the 50s measured before CCCP addition.

### Quantification of flux rate or time constants

In the ACMA-fluorescence-based K^+^ flux assay, individual fluorescence quenching curves (after CCCP addition) were fitted with a two-phase exponential equation *y*= *y*0 + *A*1 × *e*^-(*x*-*x*0)/*τfast*^*+A*2 × *e*^-(*x*-*x*0)/*τslow*^ using the Origin software (OriginLab), where A1 and A2 are amplitudes, x0 is the initial fit point, y0 corresponds to the plateau and τ is the time constant. The first point measured immediately after CCCP addition was not considered in this fit due to uncertainty in the fluorescence measurement after compound addition. Flux rate constants were determined by inverting the respective time constants (rate constant= 1/τ).

In the Pyranine-fluorescence-based K^+^ flux assay, we focused on the initial fast phase of the individual fluorescence quenching curves and estimated time constants by fitting the first 50 s after CCCP addition (50-100 s) with a one-phase exponential equation *y*= *y*0 + *A* × *e*^-(*x*-*x*0)/*τ*^ using the Origin software (OriginLab), where A is amplitude, x0 is the initial fit point, y0 corresponds to the plateau and τ is the time constant. For some of these curves, in which the fluorescence quenching change is very small over the fitted 50 s (less than 0.1 change in NF), fitting proved challenging due to a high variability of time constant values determined between similar curves. We addressed this issue by fixing y0 at the fluorescence value determined at 150 s (100 s after CCCP addition). We chose 150 s since it seems an acceptable point until which the fast phase can still be considered. By doing so, the determined time constants for these curves agree well with the measured fluxes shown on the curves. In the c-di-AMP titration assays, y0 was fixed for c-di-AMP concentrations equal or higher than 3.5 µM. Furthermore, it is important to point out that fixing y0 at 150 s does not greatly impact the time constants for the faster curves (ex for KtrA-ATP addition, time constants without and with y0 fixing are: (25.0, 22.1, 16.5, 17.8) and (20.5, 21.6, 19.8, 18.6), respectively for 4 replicas. Therefore, the time constant values determined using this method can be compared with the ones determined without fixing y0. Flux rate constants were determined by inverting the respective time constants (rate constant= 1/τ).

### TK2420 complementation assay

The complementation assay with the E. coli TK2420 strain was performed as previously described with modifications (13, 27). Wild-type KtrB and KtrA (wild type or mutants) were cloned into a dicistronic constitutive expression plasmid (pKCe). Empty pKCe plasmid was used as negative control. Transformed cells were plated on LBK (Luria–Bertani broth where NaCl is replaced by KCl) agar plates. Three individual colonies for each protein version were picked and grown separately in 5 mL of minimal media containing 30 mM K^+^, overnight at 37 □C. A 5 mL aliquot of the overnight cultures was then used to inoculate 5 mL of minimal media prepared with different K^+^ concentrations (0.1, 0.3, 1, 2, 6, 10, 30 or 115 mM) and incubated at 37 □C. Optical density at 595 nm was measured after 16 hr.

### Construction of *Bacillus subtilis* strains and plasmids

Strains and plasmids used in this study are listed in the Table S1. *E. coli* XL1-Blue was used for gene cloning.

For gene deletion in B. *subtilis* JH642, strains 168Δ*ktrA* (Δ*ktrA::kan*; BKK31090 and Δ*ktrA::erm*; BKE31090), 168Δ*ktrB* (Δ*ktrB::kan*; BKK31100), 168Δ*ktrC* (Δ*ktrC::erm*; BKE14510), 168Δ*ktrD* (Δ*ktrD::erm*: BKE13500) and 168Δ*kimA* (Δ*kimA::kan*; BKK04320) that have an antibiotic resistance cassette instead of the gene, were ordered from the Bacillus Genetic Stock Center and their genomic DNA was isolated using the GeneJET Genomic DNA Purification Kit. The antibiotic resistance cassette together with genome flanking regions (∼600 bp) were amplified and PCR products were purified using the NZYGelpure kit (NZYTech, Lisbon, Portugal). These PCR fragments were transformed into *B. subtilis*, selecting for the appropriate antibiotic resistance. Gene disruptions were confirmed by DNA sequence analysis (Eurofins, Ebersberg, Germany). The single mutant JH642 Δ*ktrA* was transformed with the PCR products of Δ*ktrC::erm* in order to obtain the double mutant Δ*ktrA*Δ*ktrC*. The single mutant Δ*ktrD* was transformed with the PCR product Δ*ktrB::kan*, in order to obtain the double mutant Δ*ktrD*Δ*ktrB*. To generate the mutant strains of 168, the lab strain 168 was first transformed with the PCR product Δ*kimA::kan* and subsequently with Δ*ktrC::erm* or Δ*ktrA::erm* in order to obtain double mutants Δ*kimA*Δ*ktrC* or Δ*kimA*Δ*ktrA*. All strains were selected using plates with kanamycin and erythromycin.

Integration of the *ktrAB* operon at *amyE* involved the cloning of two separate parts of the operon due to an apparent toxic effect of the full operon in *E. coli*. A fragment spanning part of the riboswitch together with the *ktrAB* genes was PCR amplified using JH642 genomic DNA. The purified PCR product was ligated to pDG364, giving rise to the plasmid pDG364-*ktrAB*_1. The promoter, the riboswitch and part of the *ktrA* gene were then PCR amplified using JH642 genomic DNA. The purified PCR product was ligated to pDG1730, giving rise to the plasmid pDG1730-*ktrAB*_2. The complementation of the strains with the *ktrAB* operon was done by transforming first with pDG364-*ktrAB*_1 selecting for chloramphenicol resistance and subsequently with the pDG1730-*ktrAB*_2 selecting for spectinomycin resistance. Complementation of the Δ*ktrA*Δ*ktrC* strain with *ktrA* inserted in the *amyE* locus under its own promoter was achieved by integrating the *ktrAB* operon as above but now with two stop codons in the *ktrB* gene, at the codon positions for amino acids T59 and A64.

The *ktrC* operon was amplified using JH642 genomic DNA. The purified PCR product was cloned in pDG364. To inactivate the proteins encoded by the other genes in the operon (*kinC* and *ykqA*), the H224A mutation was introduced in KinC, and four stop codons replaced codons for amino acids L5, F6, Y130 and F131 in YkqA, giving rise to plasmid pDG364-*ktrC\**.

For cloning the *ktrC* and *ktrA* genes in pTH1xp, the plasmid backbone and the genes were PCR amplified before joining the resulting fragments by isothermal DNA assembly (28). For generation of the KtrA/KtrC chimera, the N-terminal of *ktrA* was PCR amplified up to the codon for amino acid 125 and the C-terminal of *ktrC* was PCR amplified from the codon for amino acid 122 onward. The PCR products were purified and linked by cross-over PCR. The resulted PCR product was cloned into pTH1xp by isothermal DNA assembly. All cloned DNA fragments and introduced mutations were verified by sequencing.

### Culture conditions for analysis of *B. subtilis* mutants

*E. coli* and *B. subtilis* strains were routinely grown in LB-K^+^ agar plates (10 g/L of tryptone, 5 g/L yeast extract and 5 g/L potassium chloride) at 37ºC with the exception of Δ*ktrA*Δ*ktrC*-C^R169A^ strain that was selected and grown in LB. For growth experiments, *B. subtilis* was grown in Spizizen’s minimal medium (SMM). The minimal medium contains 2 g/L (NH_4_)_2_SO_4_, 1 g/L sodium citrate, 0.2 g/L MgSO_4_*7H_2_O, 0.5% glucose, 50 mg/L tryptophan and 50 mg/L phenylalanine. The phosphate salts (K_2_HPO_4_, KH_2_PO_4_, Na_2_HPO_4_ and NaH_2_PO_4_) were varied as described before (11) in order to keep a fixed pH (ratio [HPO_4_^2-^]/[H_2_PO _4_^-^] = 2), constant osmolarity ([K_2_ HPO_4_] + [Na _2_HPO4] + [KH_2_ PO_4_] + [NaH PO4] = 90 mM) and a fixed total concentration of 150 mM of monovalent cations Na^+^ and K^+^ in all solutions while varying the concentration of K^+^. Finally, 100-fold concentrated trace elements stock solution (12.5 g/L MgCl_2_*6H_2_O, 0.73 g/L CaCl_2_*2H_2_O, 1.35 g/L FeCl_2_*6H_2_O, 0.1 g/L MnCl_2_*4H_2_O, 0.17 g/L ZnCl_2_, 0.043 g/L CuCl_2_*2H_2_O, 0.03 g/L CoCl_2_ and 0.06 g/L NaMoO_4_*2H_2_O) was added at 10 mL/L.

For the potassium requirement experiments, a pre-culture of 5 mL of SMM with 40 mM K^+^ was inoculated from a LB-K^+^ agar plate and grown for about 9h at 37 ºC, 180 rpm. 5 mL of SMM with different K^+^ concentrations were inoculated with 5 µL (1000x dilution) of pre-culture. Test cultures were grown for approximately 19h at 37 ºC, 200 rpm and optical density (OD) was measured at 600 nm.

For the osmotic shock experiments, a pre-culture of 5 mL of SMM with 2 mM or 30 mM of K^+^ was inoculated from an agar plate and grown for about 9h at 37 ºC; 180 rpm. 3 mL of SMM test cultures with 2 mM or 30 mM of K^+^ were inoculated to an initial OD of approximately 0.1 and grown until ∼0.4. At this point NaCl (to a final concentration of 0.6 M) from a 5M stock solution was added to the cultures and the OD_600_ was measured 6h after.

Kanamycin (7.5 µg/mL), erythromycin (0.5 µg/mL), chloramphenicol (7 µg/mL) and spectinomycin (100 µg/mL) were added to the growth media as needed. Xylose for induction of protein expression was added in growth media from moment of inoculation.

### Analysis of the cyclic dinucleotide pools

The concentration of c-di-AMP in *B. subtilis* cells was determined by a LC-MS/MS method essentially as described previously (29). Briefly, *B. subtilis* cells (20 mL) were grown in Spizizen minimal medium at 37°C to an OD600 of 1.0. Samples (10 mL) were centrifuged (0°C, 20,800 x g), shock frozen in liquid nitrogen and stored at -80°C. This sample was used for c-di-AMP quantification. Two additional aliquots (1 mL each) were harvested for total protein determination. c-di-AMP extraction was performed as described. After addition of the internal standard [13C,15N]c-di-AMP, part of the extracts was analyzed by LC-MS/MS.

### Quantification of cyclic dinucleotides by MS/MS

The chromatographic separation was performed on a Series 200 HPLC system (PerkinElmer Life Sciences) as described previously (29). The analyte detection was performed on an API 3000 triple quadrupole mass spectrometer equipped with an electrospray ionization source (AB SCIEX) using selected reaction monitoring (SRM) analysis in positive ionization mode. The SRM transitions labeled as “quantifier” were used to quantify the compound of interest, whereas “identifier” SRM transitions were monitored as confirmatory signals. The quantifier SRM transitions were most intense and were therefore used for quantification.

## Results

### KtrC and KtrA are not physiologically redundant

To evaluate the physiological roles of KtrA and KtrC we generated single and double knockouts of all ktr genes in the *B. subtilis* JH642 strain. This strains, lacks among other genes, *kimA* that encodes the other K^+^ importer in *B. subtilis*, making it highly dependent on Ktr channels (24). Two *in vivo* assays were used to evaluate Ktr function in the deletion mutants (11). The K^+^ requirement assay reports K^+^ import activity during cell growth, while the salinity adaptation assay measures the function of Ktr channels in the adaptation mechanism to a hyperosmotic shock with NaCl, which involves a rapid uptake of K^+^ (8).

First, our results confirm the previously described roles of KtrB and KtrD (11). In particular, while the parental strain grew well with as little as 0.3 mM K^+^, the ∆*ktrB* strain requires 2 mM K^+^ for growth (Figure 1a) and does not adapt when salinity is increased (Figure 1b), confirming the idea that KtrB is an efficient K^+^ importer that has a role in hyperosmotic adaptation. The Δ*ktrB* phenotype was reversed when the *ktrAB* operon was inserted in the *amyE* locus, behaving like the parental JH642 strain (Figure S1a and S1b). In contrast, the ∆*ktrD* strain resembles the parental strain by growing in low K^+^ concentrations and adapting to hyperosmotic shock (Figure 1a and 1b). However, the double mutant Δ*ktrD*Δ*ktrB* requires at least 30 mM K^+^ to grow well (Figure 1c). Comparison with the results for the Δ*ktrB* mutant confirms that KtrB and KtrD function as K^+^ importers but in different ranges of external K^+^ concentrations, as previously proposed (11). KtrB functions in low K^+^ concentrations and KtrD has a role at high concentrations.

**Figure 1:**
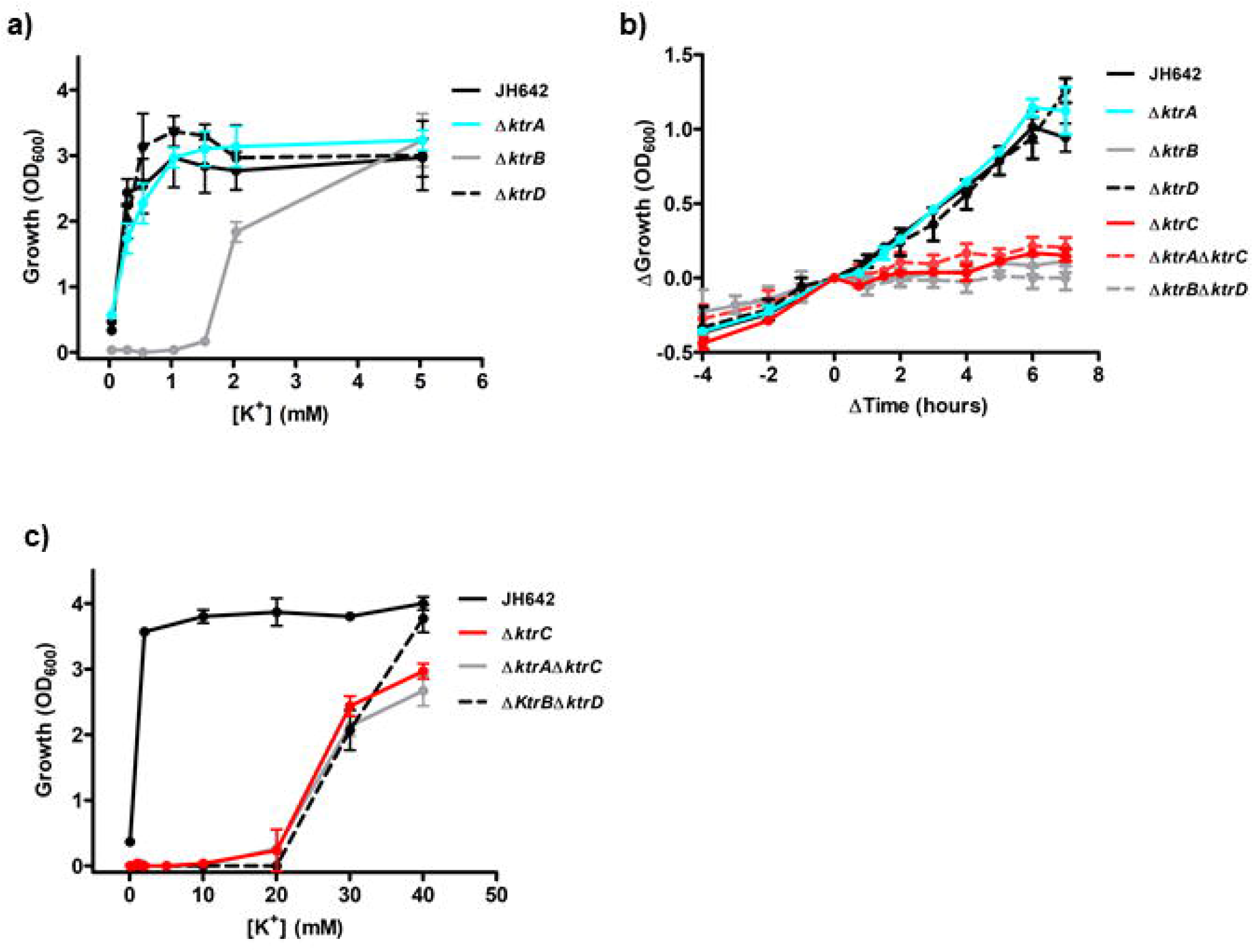
KtrC is the dominant RCK protein in *B. subtilis*. **a)** K^+^ requirement assays for *B. subtilis* JH642 strain and deletions mutant strains Δ*ktrA*, Δ*ktrB* and Δ*ktrD*. **b)** Salinity adaptation assay for *B. subtilis* JH642 strain and its deletions mutant strains Δ*ktrA*, Δ*ktrB*, Δ*ktrD*, Δ*ktrC*, Δ*ktrA*Δ*ktrC* and Δ*ktrB*Δ*ktrD*. **c)** K^+^ requirement assays for *B. subtilis* JH642 strain and deletions mutant strains Δ*ktrC*, Δ*ktrA*Δ*ktrC* and Δ*ktrB*Δ*ktrD*. Optical density at 600 nm was monitored after overnight growth in K^+^ requirement assay or over time in media with either 2 mM (JH642, Δ*ktrA*, Δ*ktrD*, Δ*ktrA*) or 40 mM K^+^ (Δ*ktrB*, Δ*ktrC*, Δ*ktrA*Δ*ktrC* and Δ*ktrB*Δ*ktrD*) in salinity adaptation assay. Hyper-osmotic shock was done at OD_600nm_ 0.3-0.5 with addition of 600 mM NaCl. For direct comparison curves from salinity assay were translated so that shock occurs at time 0 and OD_600nm =_ 0. Mean ± SD for n=3.

In contrast, the phenotypes obtained by deleting *ktrA* or *ktrC* were unexpected. The Δ*ktrA* strain grows like the parental JH642 strain in either assay (Figure 1a and 1b), displaying optimal growth in 0.3 mM K^+^ and adapting well to increased salinity. The lack of an obvious growth defect was surprising given its described close association with KtrB but it could be explained if KtrA and KtrC are redundant in the activation of KtrB, as previously highlighted by us (20). Contradicting this expectation, the Δ*ktrC* phenotype is very severe (Figure 1b and 1c). The Δ*ktrC* strain requires 30 mM K^+^ for growth, a concentration that indicates a lack of functioning dedicated K^+^ importers (compare with phenotype of Δ*ktrD*Δ*ktrB* strain, which is missing both Ktr membrane proteins), and does not recover from salinity shock. Importantly, expression of the KtrC protein under its promoter in the *amyE* locus rescued the growth phenotype of the double-deletion mutant Δ*ktrA*Δ*ktrC*, in the K^+^ requirement and salinity adaptation assays (Figure S1c and S1d). In contrast, expression of KtrA from the *amyE* locus did not rescue growth in the K^+^ requirement assay (Figure S1c). KtrA partially rescued growth in the salinity adaptation assay when expressed from the *amyE* locus, although still not as well as KtrC (Figure S1d). Note that if KtrC were to assemble and activate only KtrD then 1) deletion of *ktrC* should not be equivalent to deletion of both membrane proteins and 2) the *ktrC* complementation phenotype in the Δ*ktrA*Δ*ktrC* mutant strain should be similar to that of the Δ*ktrB* strain.

To confirm the apparent predominant role of KtrC over KtrA in a legacy laboratory strain, we deleted the genes from the *B. subtilis* strain 168. We first deleted *kimA*, the other major K^+^ importer in *B. subtilis* and in this background we generated the strains ∆*kimA*∆*ktrA* and ∆*kimA*∆*ktrC*. These strains were tested in the K^+^ requirement assay (Figure S1e). The lack of the gene encoding for KimA did not have a noticeable effect and the strain grew in all K^+^ concentrations as expected from the presence of active Ktr channels. However, in the double mutants we saw two distinct results; deletion of *ktrA* did not alter the growth properties relative to the *kimA* deletion mutant at concentrations of 0.5 mM and higher, while deletion of *ktrC* only allowed growth in 40 mM K^+^. These results mirror our findings with the JH642 strain.

Our results show that in the cell, K^+^ transport mediated by Ktr cation channels is KtrC-dependent. KtrC is essential not just for KtrD activation, as previously proposed, but also for the function of the major *B. subtilis* Ktr membrane protein, KtrB. In turn, KtrA is not sufficient to activate K^+^ import by Ktr channels, although it might have a role in adaptation to hyperosmotic shock.

### KtrC mediates c-di-AMP inactivation

To understand the distinct physiological role of KtrC and KtrA, we analyzed their functional properties *in vitro*. For this, we assembled the two RCK proteins with the same membrane protein, KtrB, and verified that KtrCB-ATP has a similar elution profile to KtrAB in size exclusion chromatography (Figure S1a). We then used a liposome flux-assay (21) with the H^+^-sensitive fluorescent dye ACMA to demonstrate that KtrC-ATP activates KtrB as well as KtrA-ATP (Figure S2b and S2c). All ion flux experiments were performed in the presence of activating 100 µM ATP and 1 mM Mg^2+^ (21). Liposomes reconstituted with KtrCB-ATP and KtrAB-ATP showed similar flux curves, with fast rate constants of 0.056 ±0.003 and 0.053 ±0.001 s^-1^, respectively. In order to exclude potential differences in the liposome reconstitution efficiencies of KtrCB and KtrAB, we also prepared liposomes with KtrB alone and assembled the complex by adding an excess of KtrC or KtrA to the outside solution (up to a maximum of 120 nM of RCK protein) (Figure S2d). This approach ensures that we have the same amount of membrane protein in the assay, allowing a direct comparison of the activation of KtrB by KtrC or KtrA. Again, the flux curves are very similar with fast rate constants of 0.053±0.011 and 0.051±0.009 s^-1^ for KtrC and KtrA, respectively (Figure 2a).

**Figure 2:**
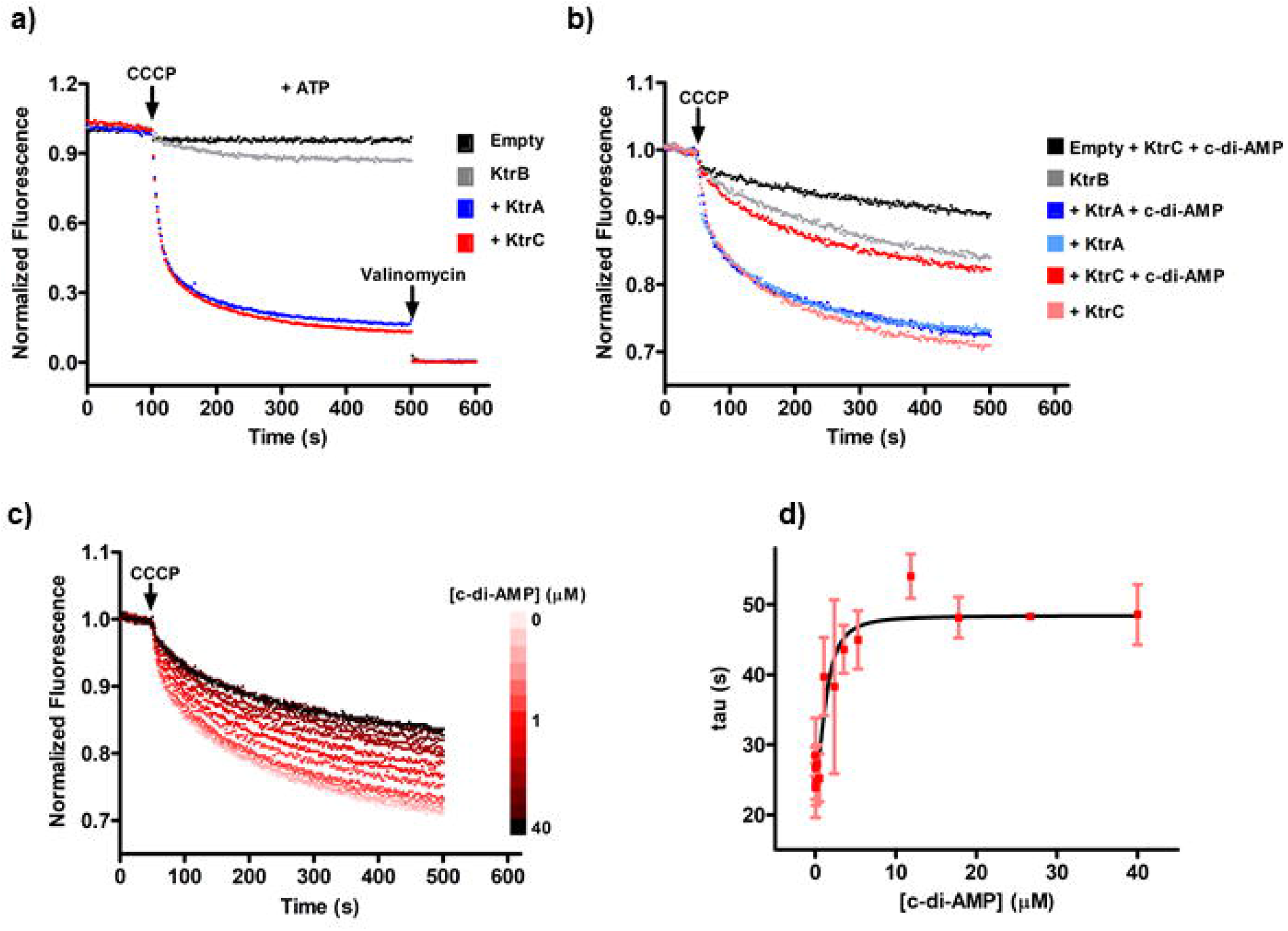
c-di-AMP inhibits Ktr channels assembled with KtrC but not KtrA. **a)** ACMA fluorescence flux curves for Ktr channels assembled with addition of KtrA or KtrC to KtrB-reconstituted liposomes. **b)** Pyranine fluorescence flux curves for Ktr channels assembled with addition of KtrA or KtrC to KtrB-reconstituted liposomes, with and without 20 µM c-di-AMP. **c)** Example of pyranine fluorescence flux curves resulting from titration of KtrCB with increasing concentrations of c-di-AMP. **d)** Plot of time constants extracted from c-di-AMP titration curves as a function of c-di-AMP concentration. Data were fitted with Hill equation with the following parameters, Ki= 1.4 ±0.5 μM and Hill coefficient n= 1.9 ±0.7. All assays were performed in the presence of ATP and Mg^2+^. Mean ±SD for n=3.

As we did not observe differences in the activation properties of the two RCK proteins, we wondered if the 100-fold difference in the constant of dissociation of c-di-AMP binding to KtrC or KtrA (20) could underlie a distinct functional effect. However, c-di-AMP interfered with the ACMA flux-assay, probably by interacting with the dye. Therefore, we setup a similar assay but using pyranine, a water-soluble H^+^-sensitive fluorescent dye that is encapsulated in the liposome and is not exposed to the externally added dinucleotide (30) (Figure S2e). Using this assay, we could demonstrate that 20 µM c-di-AMP inhibits KtrCB-ATP but not KtrAB-ATP (Figure 2b). We performed a functional titration with c-di-AMP and determined the rate constants from single exponentials fitted to the first 50 sec of each curve. A plot of flux time constants as a function of dinucleotide concentration and fitting the Hill equation showed that the c-di-AMP concentration at 50% inhibition (Ki) is 1.4 µM with a Hill coefficient (n) of 1.9. In contrast, flux curves measured with KtrAB-ATP in the presence of 50 µM c-di-AMP (Figure S2f) do not show a difference in the rate constant (0.052 ±0.003 s^-1^) relative to curves measured in the absence of dinucleotide (0.056 ±0.006 s^-1^). However, comparison of averaged curves in these two conditions shows a change in the curve shape beyond 200 seconds which suggests that KtrA regulated channels can be inhibited at very high c-di-AMP, in agreement with the difference in the c-di-AMP binding affinity of the isolated RCK proteins (20).

Overall, our *in vitro* results demonstrate that KtrC and KtrA activate KtrB equally well. Importantly, the two RCK proteins are distinct in their ability to promote channel inactivation by c-di-AMP; while KtrC mediates inactivation at low micromolar concentrations of c-di-AMP, KtrA is very insensitive to the dinucleotide.

### KtrA and KtrC form heteromers that are inactivated by c-di-AMP

Surprisingly, we found that KtrA and KtrC assemble as heteromeric rings when we co-expressed Strep-tagged KtrA and His-tagged KtrC in *E. coli*. Pulling-down Strep-KtrA with Streptactin beads revealed the presence of the two proteins (Figure 3a). The same happened when His-KtrC was pulled-down with Co^2+^ beads (Figure S3a). Control experiments showed that Strep-KtrA alone or His-KtrC alone are not pulled-down by Co^2+^ or Streptactin beads, respectively (Figure 3a and S3a). The high amino acid sequence similarity between KtrA and KtrC (55% sequence identity) explains the formation of these heteromers. Moreover, *in-vitro* mixing of purified Strep-KtrA and untagged KtrC, with either bound ATP or ADP and with or without c-di-AMP, followed by pull-down with Streptactin, revealed that untagged KtrC associated with KtrA in all conditions (Figure 3b and S3b). This association is slow, taking hours for completion, indicating that it does not result from a non-specific interaction between KtrA and KtrC and is instead consistent with a process involving conformational changes. Size-exclusion chromatography analysis of the Strep-KtrA/KtrC complex (_Strep_KtrA/C), formed by mixing the two proteins together and purified with Streptactin beads, showed the complex eluting at the same volume as octameric KtrA (Figure S3c). It is worthwhile pointing out that it is very likely that this preparation is composed by a heterogeneous population of octameric rings with different ratios of KtrA and KtrC subunits. Incubation of _Strep_KtrA/C with KtrB gives rise to a size-exclusion peak that contains all three proteins (Figure S3c and S3d) and has an elution volume similar to KtrAB or KtrCB (cf. Figure S3c with S2a). SDS-PAGE analysis of _Strep_KtrA/CB complex reconstituted in liposomes also showed the presence of the three proteins (Figure S3d).

**Figure 3:**
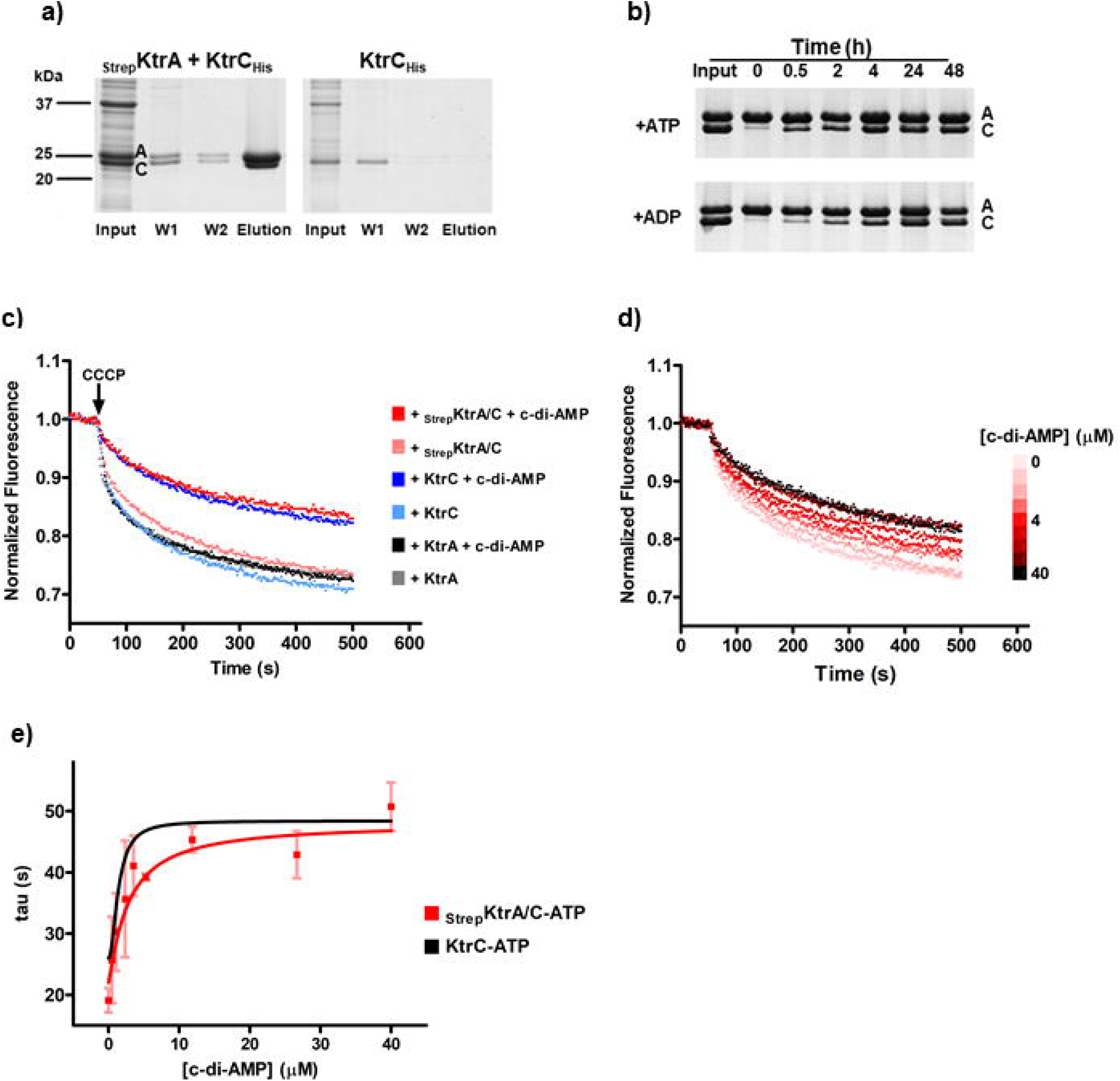
Assembly and function of KtrA/C heteromeric RCK proteins. **a**) SDS-PAGE analysis of fractions collected from a pulldown experiment with Streptactin beads. Pulldown was performed with a lysate of *E. coli* cells (Left) co-expressing N-terminally Strep-tagged KtrA (_Strep_KtrA) and C-terminally His-tagged KtrC (KtrC_His_) or (Right) expressing KtrC_His_ alone, showing that KtrC_His_ is present in the elution fractions only in the co-expression experiment. Lysare (input), washes (W1,W2), elution with desthiobiotin. **b**) SDS-PAGE analysis of eluted fractions from pulldown experiments performed with samples collected at various time points from a mixture of _Strep_KtrA and KtrC (independently purified) prepared and incubated in the presence of ATP (top) and ADP (bottom) **c**) Pyranine fluorescence flux curves for Ktr channels assembled by addition of KtrA, KtrC or heteromeric _Strep_KtrA/C to KtrB-reconstituted liposomes in the presence of ATP and Mg^2+^ with and without 20 µM c-di-AMP. d) Example of pyranine fluorescence flux curves resulting from titration of assembled KtrA/CB with increasing concentrations of c-di-AMP. **e)** Plot of time constants extracted from curves in d) as a function of c-di-AMP concentration. Data were fitted with Hill equation (red curve) with the following parameters, Ki= 2.8 ±0.6 μM and Hill coefficient n= 1.1 ±0.4. For comparison, we show the fitted curve (black line) to the c-di-AMP titration of KtrCB (from Figure 2d). Mean ±SD from n=3.

Interestingly, the ability to form hetero-oligomers is not restricted to the two RCK proteins from *B. subtilis* and is present in RCK protein pairs from other species. In particular, in *Streptococcus mutans* and *Salinicoccus halodurans*, which have two RCK proteins but different genomic organizations (Figure S4a). Streptactin pull-down experiments of co-expressed RCK proteins in *E. coli* revealed the two proteins in the elution fractions, demonstrating association (Figure S4b).

We also considered whether KtrB and KtrD could form hetero-dimers when recombinantly co-expressed. However, pull-down experiments after detergent solubilization showed no evidence for formation of KtrB/KtrD dimers (Figure S5).

We wondered if KtrA/C heteromeric rings are functional and sensitive to c-di-AMP inhibition. For this we optimized the *in vitro* conditions for assembly of the _Strep_KtrA/CB complex to get similar amounts of the two proteins in the pulled-down complex and analyzed the functional properties of the complex by adding it to KtrB liposomes. The resulting flux curves are similar to KtrAB and KtrCB curves (Figure 3c). Importantly, addition of 20 µM c-di-AMP inhibits flux as completely as for KtrCB liposomes. If the _Strep_KtrA/CB protein preparation was composed by a mixture of _Strep_KtrAB and KtrCB complexes then we would expect to observe only partial flux inhibition since channel complexes formed with KtrA are insensitive to this concentration of c-di-AMP. Complete inhibition reinforces the idea that we are analyzing the function of a Ktr channel assembled with 3 proteins. c-di-AMP titration using the liposome flux assay revealed a concentration-dependence that is unchanged relative to the KtrCB (Ki 2.8 µM) (Figures 3d and 3e). However, the titration curve rises less steeply, as reflected in a Hill coefficient that is close to ∼1 (n= 1.1), possibly reflecting a change in the number of c-di-AMP binding sites or in the c-di-AMP induced mechanism of inhibition.

These results show that the regulatory network formed by bacterial RCK proteins is potentially more complex than previously thought. It is likely that in some bacterial species we need to also consider hetero-octamers assembled from different RCK proteins. Importantly, in *B. subtilis* the presence of KtrC in the heteromeric rings confers sensitivity to inhibition by c-di-AMP.

### c-di-AMP sensitivity explains the dominant role of KtrC

To explore how the sensitivity of KtrC to c-di-AMP affects its physiological function and how it might relate to its dominant role, we evaluated the impact of mutations in the c-di-AMP binding site.

We first analyzed how complementation of the JH642 Δ*ktrA* Δ*ktrC* strain (Δ*ktrA*Δ*ktrC*) by the *ktrC* operon inserted in the *amyE* locus is affected by a mutation that reduces c-di-AMP binding. For this we mutated to alanine the conserved arginine (R169 in KtrC) that has been shown to be essential for c-di-AMP binding to the C-terminal subdomain of RCK proteins (22, 31). We confirmed that the KtrC_R169A_ mutant is functional and activates KtrB with the *E. coli* strain TK2420 complementation assay (Figure S6a); this strain requires expression of an active K^+^ transporter or channel to grow with less than 30 mM K^+^. We also tested the function of the KtrC_R169A_ mutant protein in the liposome flux assay (Figure S6b). Addition of KtrC_R169A_-ATP to the outside of KtrB-containing liposomes gives rise to flux curves that are indistinguishable from wild type KtrC-ATP. As expected, external addition of 20 μM c-di-AMP does not inhibit flux, confirming the effect of the R169A mutation in c-di-AMP binding. We then compared the physiological impact of wild type and R169A *ktrC* operon. As above, wild type KtrC expressed from its own promoter rescued the Δ*ktrA*Δ*ktrC* strain phenotype at all K^+^ concentrations tested in the K^+^ requirement assay (Figure 4a). In contrast, the strain with the mutant *ktrC* grew poorly in most conditions with the exception of those with lowest amounts of K^+^ (0.2 and 0.5 mM K^+^) where OD was ∼1. Strikingly, these are also the conditions expected to have the lowest amounts of c-di-AMP (24). These results demonstrate that the physiological function of KtrC is highly dependent on c-di-AMP binding and inhibition. However, we worried about possible confounding effects imposed by native regulation of transcription and/or translation of the *ktrA* and *ktrC* genes. Therefore, we expressed these proteins and their mutants under a xylose-inducible promoter from the replicative plasmid pTH1xp (32) in the Δ*ktrA*Δ*ktrC* strain.

**Figure 4:**
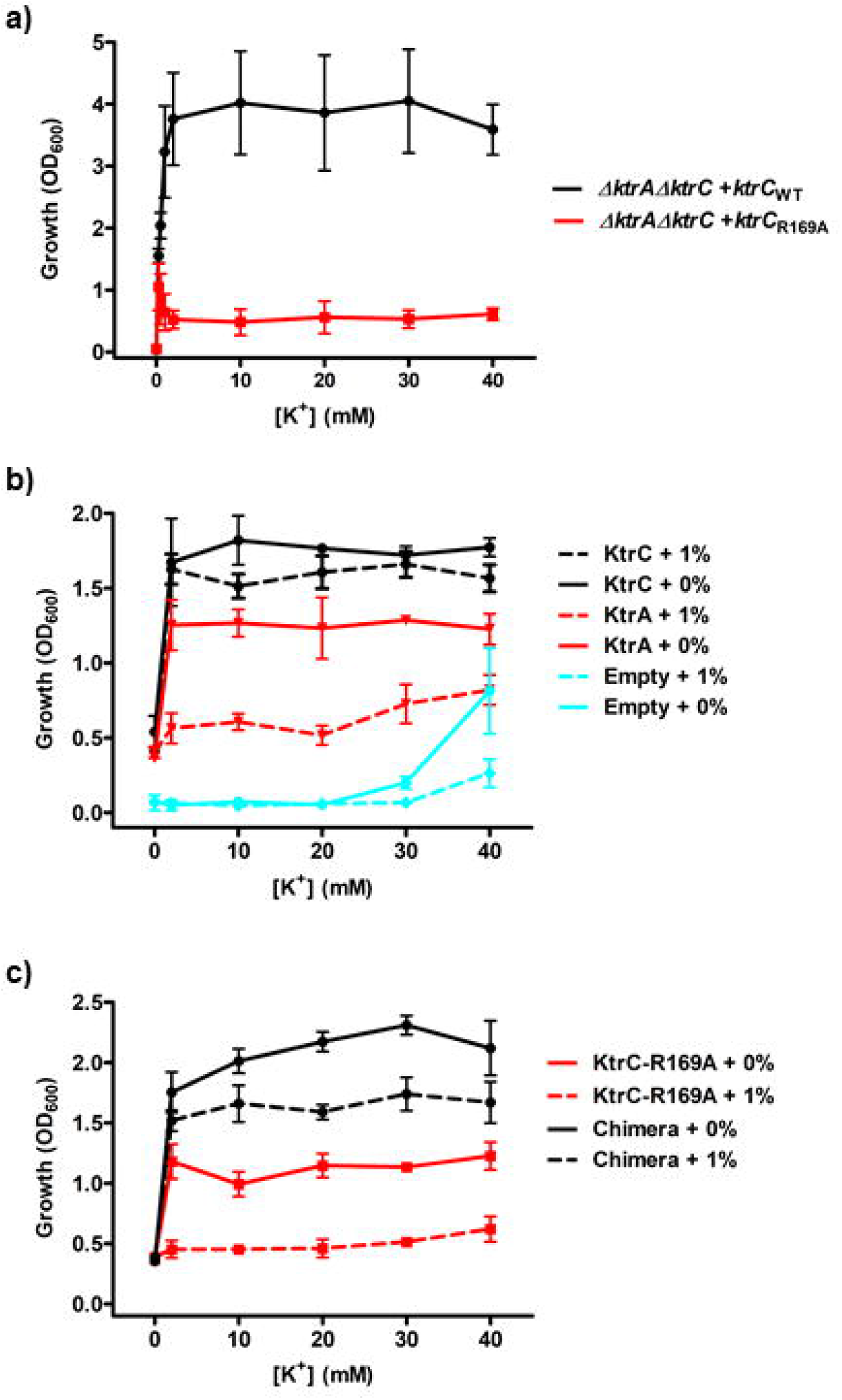
Intact c-di-AMP site determines toxicity of over-expressed RKC protein. K^+^ requirement assays for **a)** *B. subtilis* JH642 Δ*ktrA*Δ*ktrC* strain complemented with wild type *ktrC* or mutant *ktrC*_R169A_ from *ktrC* operon integrated in the *amyE* locus, **b)** JH642 ∆*ktrA*∆*ktrC* strain transformed with pTH1xp plasmid encoding KtrA or KtrC and with (1% xylose) or without induction (0% xylose) of protein expression and for **c)** JH642 ∆*ktrA*∆*ktrC* strain transformed with empty pTH1xp plasmid or encoding KtrC_R169A_ or KtrA/KtrC chimera with (1% xylose) or without induction (0% xylose) of protein expression. Overnight growths were monitored by optical density at 600 nm in media with different K^+^ concentrations which contained xylose (when appropriate) from the moment of inoculation. Mean ± SD for n=3.

We found that leaky expression in the absence of inducer (0% xylose) was sufficient for KtrA and KtrC to complement the severe growth phenotype at all K^+^ concentrations tested (Figure 4b). Although strains transformed with pTH1xp-*ktrA* show slightly lower OD_600nm_ values than those with pTH1xp-*ktrC*, these results confirm that KtrA is able to complement the growth phenotype by activating Ktr channels in the cell. In contrast, induction with 1% xylose showed very different results. Cells expressing KtrC grew as well as before but cells expressing KtrA grew poorly, revealing that the expression level of KtrA affects its physiological function most likely by assembling and activating a larger fraction of Ktr channels. Importantly, the growth of Δ*ktrA*Δ*ktrC* strain complemented with the KtrC_R169A_ mutant (pTH1xp-*ktrC*_*R169A*_) recapitulates the behavior of the strain expressing KtrA, it grew well without induction and poorly with 1% xylose (Figure 4c). Altogether, these experiments indicate a strong correlation between a lack of c-di-AMP inhibition and an inability to complement the growth phenotype at high protein expression levels.

To strengthen further this conclusion, we modified KtrA to acquire the c-di-AMP binding site of KtrC. For this, we generated a chimera protein fusing the N-terminal subdomain of KtrA, which is responsible for the formation of the regulatory ring and the interaction with the Ktr membrane protein, with the C-terminal subdomain of KtrC, which includes the c-di-AMP binding site. When co-expressed with KtrB, the chimera was functional and rescued the phenotype of *E. coli* TK2420 (Figure S6a). Importantly, the chimera behaved like KtrC when expressed from the replicative plasmid pTH1xp, with the strain growing well with and without induction, supporting the parallel between c-di-AMP inhibition and robust growth at high expression levels. We could not verify *in vitro* if the chimera is inhibited by c-di-AMP because the purified protein did not behave as well as the other RCK proteins when isolated. However, introduction of the equivalent mutation to R169A in the chimera gave rise to a phenotype that resembled KtrC_R169A_ strong complementation with 0% and poor complementation with 1% xylose (Figure S6c). This further supports the idea that the ability of the RCK protein to bind c-di-AMP and mediate c-di-AMP inhibition correlates with the complementation of the growth phenotype at all levels of protein expression. To ensure that the correlation between growth phenotypes with xylose induction and c-di-AMP is not due to unforeseen deregulation of c-di-AMP homeostasis by addition of xylose, we determined that c-di-AMP levels are unchanged by xylose addition in strains JH642(pTH1xp) and JH642 Δ*ktrA* Δ*ktrC* (pTH1xp) (Figure S7).

In summary, these experiments confirm that both KtrA and KtrC are able to activate Ktr channels in the cell. However, for KtrA this effect is dependent on its protein levels. High expression levels of KtrA, which is not sensitive to c-di-AMP, reduce growth at all K^+^ concentrations relative to low protein expression levels. The same is observed with the KtrC mutant that is not regulated by c-di-AMP. In contrast, KtrC and the chimera, both containing a wild-type c-di-AMP binding site, are able to promote strong growth at low and high expression levels.

Altogether, the data reveal that it is toxic to the cell to have a large fraction of Ktr channels that cannot be controlled by c-di-AMP. Therefore, we propose that the “physiologically dominant role” of KtrC over KtrA in the activation of Ktr channels is linked to the ability of c-di-AMP to bind to KtrC and regulate Ktr channel activity, avoiding the toxic effects of K^+^ over-accumulation.

## Discussion

Our results offer a radically different view of the physiological organization of the RCK proteins in the model organism *B. subtilis*. First, we showed that KtrC is the physiologically dominant RCK protein since deletion of KtrC is equivalent to deletion of all Ktr membrane proteins while deletion of KtrA has no impact on Ktr channel mediated K^+^ transport. This KtrC-dependent Ktr channel activity implies that KtrC, KtrA and their membrane protein partners do not just assemble according to the simple model of KtrAB and KtrCD. Instead, at least KtrCB must also be present in the cell since the function of KtrB is essential for growth in low K^+^ levels (below 2 mM). The predominance of KtrC fits well with the higher level of transcription for *ktrC* relative to *ktrA* (33) and high average number of KtrC protein molecules estimated to exist in a cell (varying between 450 to 6000 copies) (34). Moreover, we provided evidence that the universe of Ktr channels is also likely to include KtrA/CB and KtrA/CD complexes since functional RCK heteromeric assemblies are easily formed both *in vitro* and in *E. coli*. Further experiments are required to determine what is fraction of Ktr channels that in *B. subtilis* are assembled with KtrA, KtrC or KtrA/C. Moreover, we showed that the potential existence of heteromeric assemblies is not restricted to *B. subtilis* and is also possible in *Streptococcus mutans* and *Salinicoccus halodurans*. In addition, others had previously shown that two RCK proteins in *Thermatoga maritima*, with their genes in the same locus, also form hetero-oligomers (35, 36). It is therefore likely that the complex organization of Ktr channels seen in *B. subtilis* is present across different bacterial species.

Second, we showed that although both KtrC and KtrA assemble with KtrB and mediate similar K^+^ flux, only a channel assembled with KtrC (as KtrCB or KtrA/CB) is inhibited by low concentrations of c-di-AMP. In contrast, KtrAB fluxes are unchanged by addition of c-di-AMP up to 20 µM, appearing to show some inhibition at 50 µM. The range of physiological concentrations of c-di-AMP in *B. subtilis* cells remains undetermined and therefore it is difficult to judge in what conditions Ktr channels are inhibited by c-di-AMP. Nevertheless, a simple estimate based on available data (see supplemental information for details) suggests that a reasonable lower limit for c-di-AMP is 0.1 µM (in conditions where net K^+^ uptake is required) while the maximum concentration is around 30 µM, for example during the adaptation to hyperosmotic shock when net K^+^ export is required. At 0.1 µM c-di-AMP the majority of KtrCB and KtrA/CB channels in the cell is in the apo-conformation and active, favoring the import of K^+^. On the other hand, at 30 µM c-di-AMP, close to 100% of Ktr channels regulated by KtrC or KtrA/C are inactive while a large fraction of those regulated by KtrA alone remain active.

Importantly, we have demonstrated that having a large population of channels assembled with KtrA alone is a problem for the cell., Therefore, this strongly indicates that the majority of Ktr channels in the cell must be assembled and regulated by KtrC or KtrA/C, explaining the evolutionary selection of KtrC as the dominant RCK protein dedicated to K^+^ transport in *B. subtilis*. The importance of KtrC and the role of c-di-AMP in its establishment fits well with the isolation of KtrC suppressor mutants, and not of KtrA, in c-di-AMP free *B. subtilis* strains (12, 37, 38).

The modest role of KtrA in K^+^ transport revealed here does not rule out a more significant participation in other cellular processes. In particular, the link between KtrA and KtrB, through the KtrAB operon, suggest that this RCK protein is not just a genetic “archaeological remain”. In addition, our quantitative demonstration that Ktr channels are regulated by c-di-AMP raises questions about the physiological relationships between the regulation of these channels by c-di-AMP, ATP, ADP and glutamate, which has been proposed to stimulate K^+^ uptake by KtrD (12).

Altogether, our results fully support the proposal that c-di-AMP is the master regulator of the K^+^ machinery in *B. subtilis*. c-di-AMP controls this machinery at the genetic level, by regulating the transcription of *kimA* and of the *ktrAB* operon (23, 24) and at the protein level, by directly regulating the function of Ktr channels, of KimA (a K^+^/H^+^ symporter that seems to be inhibited by c-di-AMP) (37, 39) and of KhtTU (a K^+^/H^+^ antiporter that is activated by c-di-AMP) (31). We now show that the role of c-di-AMP goes beyond this direct regulation and also imposes constraints on the hierarchical control of Ktr channels by RCK proteins.

## Acknowledgements

We are grateful to Heike Bähre and Roland Seifert for c-di-AMP quantification. We acknowledge the SOLEIL and ALBA synchrotrons and the scientific platforms of i3S (Biochemical and Biophysical Technologies, Proteomics and X-ray Crystallography) for access and thank their staff for help with data collection.

This work was supported by FEDER - Fundo Europeu de Desenvolvimento Regional funds through the COMPETE 2020-Operational Programme for Competitiveness and Internationalization (POCI), Portugal 2020, and by Portuguese funds through FCT - Fundação para a Ciência e a Tecnologia/Ministério da Ciência, Tecnologia e Ensino Superior in the framework of the projects POCI-01–0145-FEDER-029863(PTDC/BIA-BQM/29863/2017), by the Fundação Luso-Americana para o Desenvolvimento through the FLAD Life Science 2020 award ‘Bacterial K^+^ transporters are potential antimicrobial targets: mechanisms of transport and regulation’, by “la Caixa” Foundation under the project code “LCF/PR/HR22/52420004” (all to J.H.M.-C.) and a grant of the Deutsche Forschungsgemeinschaft (DFG) within the Priority Program SPP1879 (Stu 214-16) (to J.S.). The funders had no role in study design, data collection, analysis and interpretation, decision to submit the work for publication, or preparation of the manuscript.

