## Supplemental Figures and Table for "c-di-AMP determines the hierarchical organization of bacterial RCK proteins"

**Estimation of c-di-AMP concentration range in B. subtilis**

It has been demonstrated that $K^+$ is a major factor in determining c-di-AMP levels, rising 2-fold for *B. subtilis* growing in a modified Spizizen minimal medium between 0.1 mM and 5 mM $K^+$ (1). An absolute quantification of c-di-AMP levels in *B. subtilis* indicated that the concentration of c-di-AMP in vegetative cells is 1.7 µM (2). These cells grew in hydrolyzed casein (CH) growth medium which is likely to contain more than 10 mM of $K^+$. We have now shown that $K_i$ for inhibition of KtrCB is ~1 µM. So, it seems safe to assume that the lower range of c-di-AMP concentration is well below 1 µM, let us say ~0.1 µM, so that a large fraction of KtrCB channels is active in low $K^+$ conditions. On the other hand, in a study where cells grew in a modified minimal media with ~10 mM $K^+$ (3) it was determined that *B. subtilis* had 190-fold more c-di-AMP than in the study with modified Spizizen minimal medium. Moreover, c-di-AMP concentration dropped 4-fold immediately after hyperosmotic shock with 1.2 M NaCl followed by a 7-fold rise during the adaptation process (3). Taking 0.1 µM as the lower limit this means that c-di-AMP can rise to ~30 µM in a condition that requires active reduction of intracellular $K^+$ through inactivation of $K^+$ importers and activation of $K^+$ exporters.

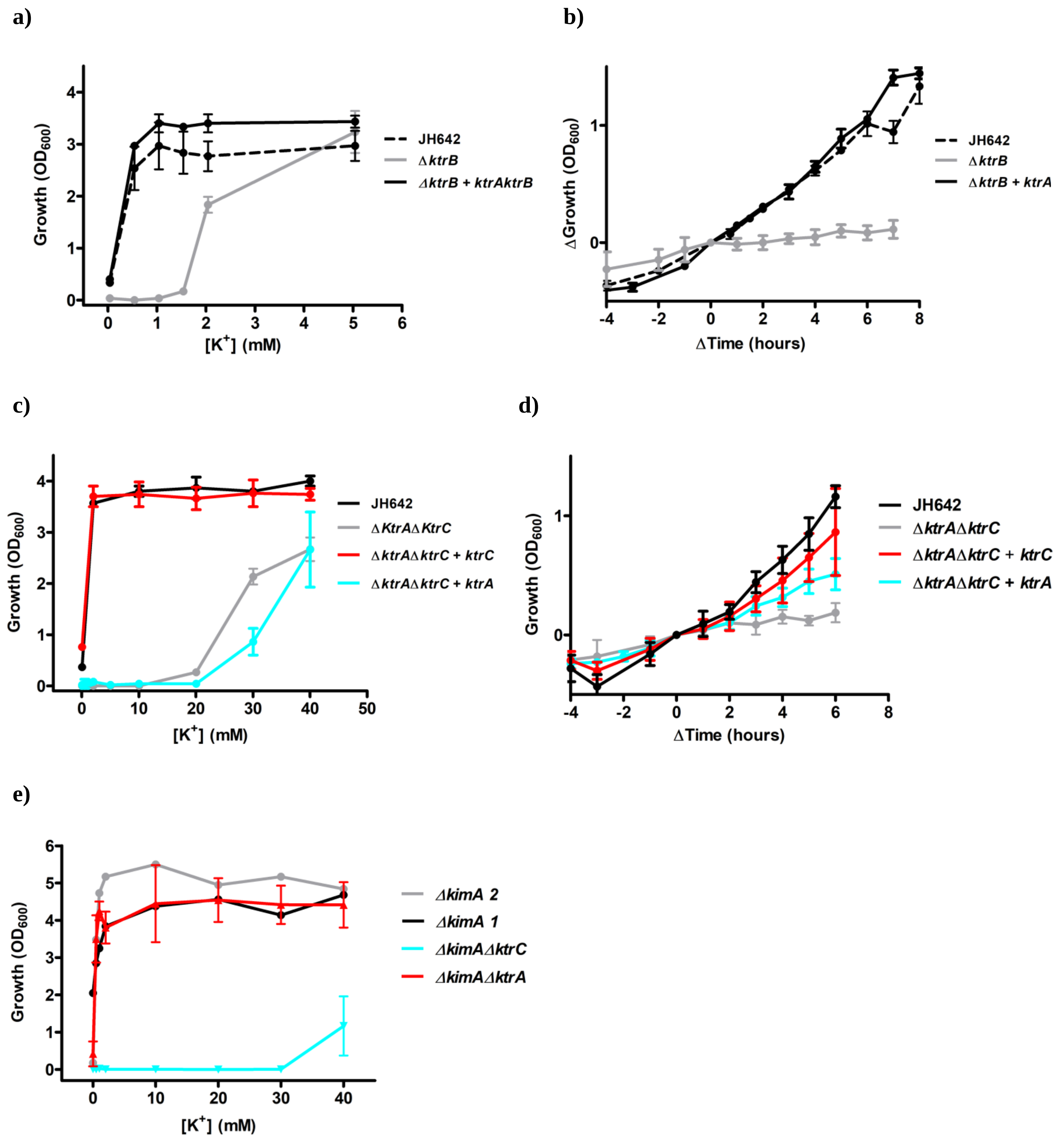


**Figure S1: Phenotype characterization of Ktr gene deletion mutants. a)** $K^+$ requirement assay for *B. subtilis* JH642 strain and mutant strains *ΔktrB* and *ΔktrB* with the *ktrAB* operon integrated in the *amyE* locus. **b)** Salinity adaptation assay for *B. subtilis* JH642 strain and mutant strains *ΔktrB* and *ΔktrB* with the *ktrAB* operon integrated in the *amyE* locus. **c)** $K^+$ requirement assay for *B. subtilis* JH642 strain and mutant strains *ΔktrAΔktrC*, *ΔktrAΔktrC* with *ktrC* or *ktrA* genes (under the control of their own promoters) integrated in the *amyE*. **d)** Salinity adaptation assay for *B. subtilis* JH642 strain and mutant strains *ΔktrAΔktrC* and *ΔktrAΔktrC* with the *ktrC* or *KtrA genes* (under the control of their own promoters) integrated in the *amyE* locus. **e)** $K^+$ requirement assay for *B. subtilis* 168 strain *ΔkimAΔktrA* and *ΔkimAΔktrC* and 2 clones of strain *ΔkimA*. For $K^+$ requirements assays, overnight growth was monitored by optical density at 600 nm in media with different $K^+$ concentrations. For salinity adaptation assays, optical density at 600 nm was monitored over time in media with either 2 mM (JH642, *ΔktrB* with *ktrAB* operon in *amyE*, and *ΔktrAΔktrC* with *ktrC gene* in *amyE*) or 40 mM $K^+$ (*ΔktrB*, *ΔktrAΔktrC* and *ΔktrAΔktrC* with *ktrA* operon in *amyE*). For salinity adaptation assays, hyper-osmotic shock with addition of 600 mM NaCl was done at $OD_{600nm}$= 0.3-0.5.  Curves were translated so that shock occurs at time 0 and $OD_{600nm}$= 0. Mean ± SD for n=3.

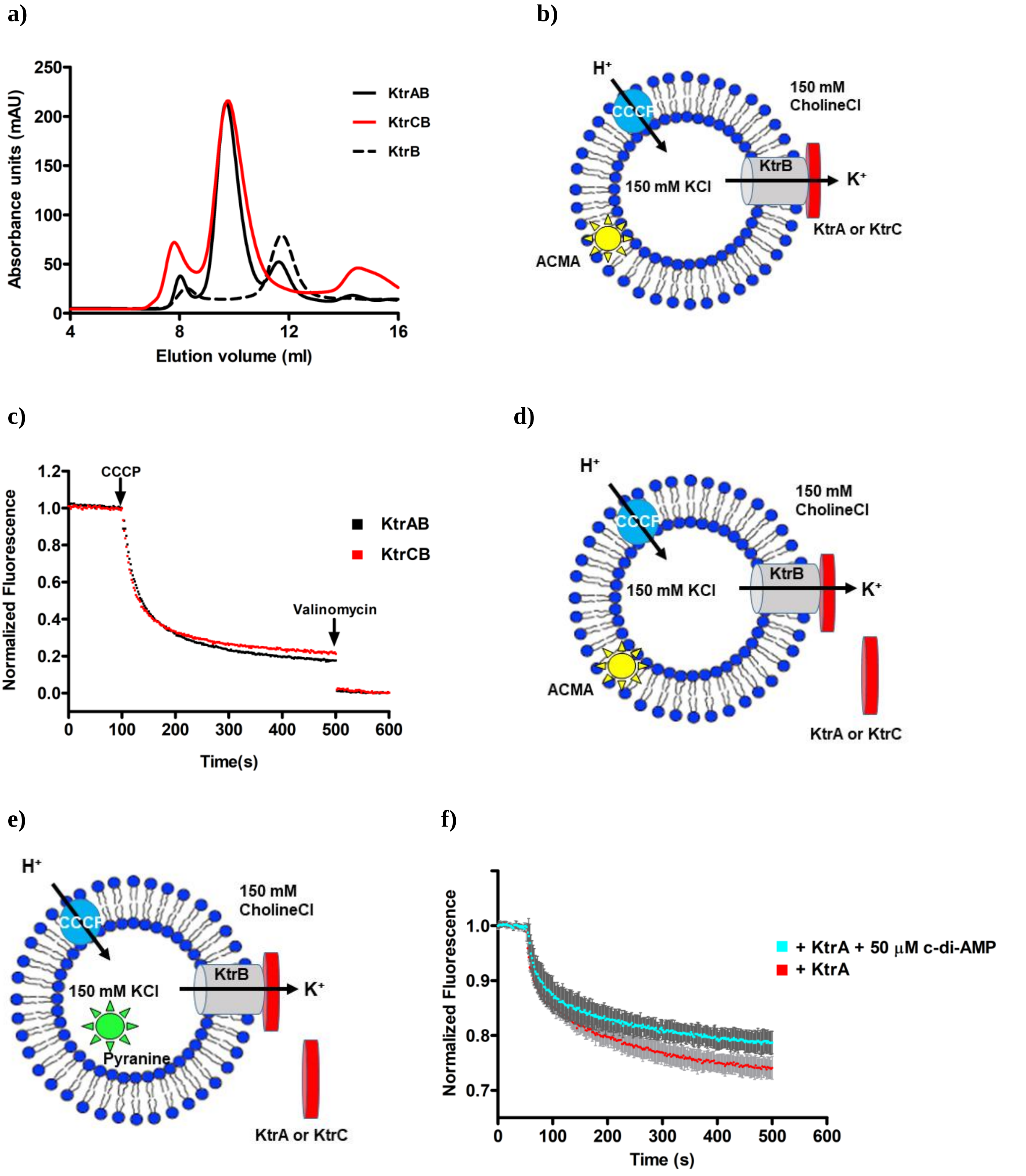


**Figure S2: Assembly and liposome reconstitution of KtrAB and KtrCB channels. a)** Comparison of size-exclusion chromatography elution profiles for KtrAB, KtrCB and KtrB. **b)** Cartoon showing configuration of ACMA liposome flux assay with fully-assembled KtrAB or KtrCB channels reconstituted in liposomes. Flux is initiated by addition of the $H^+$-ionophore CCCP and $H^+$ uptake results from the outward flux of $K^+$ from the liposome lumen down its concentration gradient. Accumulation of

$H^+$ in the liposome membrane is detected by fluorescence quenching of the fluorophore ACMA. **c)** ACMA fluorescence flux curves measured for KtrAB and KtrCB channels reconstituted in liposomes and in the presence of ATP and $Mg^{2+}$. **d)** Cartoon showing confirguration of ACMA liposome flux assay with KtrB-reconstituted liposomes and addition of KtrA or KtrC to the outside of liposome. **e)** Cartoon showing configuration of pyranine liposome flux assay monitored with KtrB-reconstituted liposomes and addition of KtrA or KtrC to the outside of liposome. Pyranine is encapsulated in the liposome and its fluorescence is quenched by accumulation of $H^+$. **f)** Comparison of averaged flux curves for KtrAB in the presence of ATP, $Mg^{2+}$ with and without 50 µM c-di-AMP. Mean ±SD for n=4.

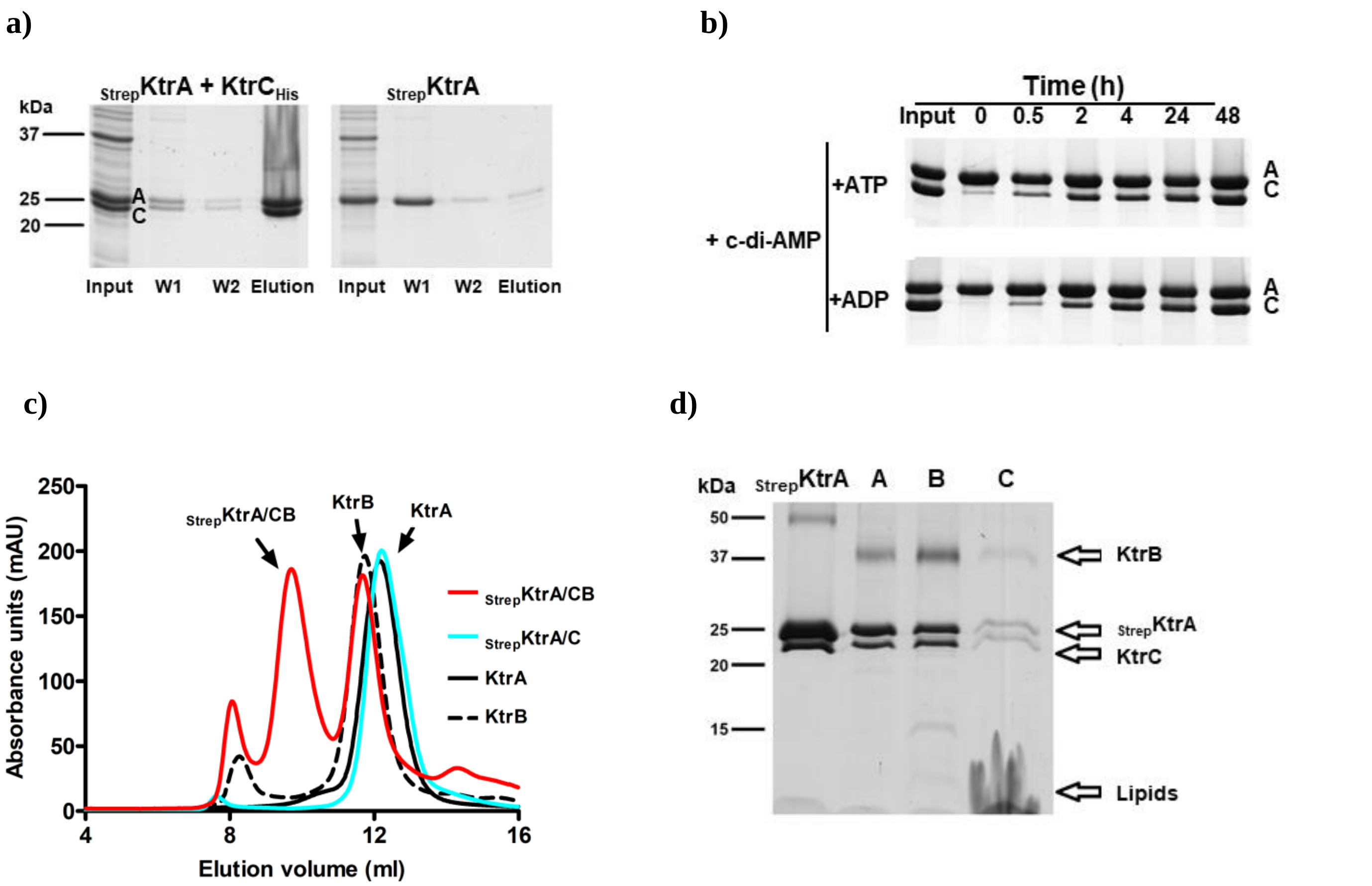


**Figure S3: KtrA/C assembly. a)** SDS-PAGE analysis of fractions collected from a pulldown experiment with Ni$^{2+}$ beads. Pulldown was performed with a lysate of *E. coli* cells (Left) co-expressing N-strep tagged KtrA ($_{Strep}$KtrA) and C-terminally His tagged KtrC (KtrC$_{His}$) or (Right) expressing $_{Strep}$KtrA alone, showing that $_{Strep}$KtrA is present in the elution fractions only in the co-expression experiment. Lysate (input), washes (W1,W2) elution with 150 mM Imidazole. **b**) SDS-PAGE analysis of eluted fractions from pulldown experiments performed with samples collected at various time points from mixture of $_{Strep}$KtrA and KtrC (independently purified) prepared and incubated in the presence of ATP (top) and ADP (bottom) with added c-di-AMP **c)** Superposition of size-exclusion chromatography elution profiles of KtrA, KtrC, $_{Strep}$KtrA/C and $_{Strep}$KtrA/CB. **d)** SDS_PAGE analysis of $_{Strep}$KtrA and KtrC mixture ($_{Strep}$KtrA/C) after purification with Streptactin beads, (A) mixture of KtrB and $_{Strep}$KtrA/C at 1:1 (w:w), (B) $_{Strep}$KtrA/CB elution peak from size-exclusion chromatography and (C) of $_{Strep}$KtrA/CB reconstituted in proteoliposomes.

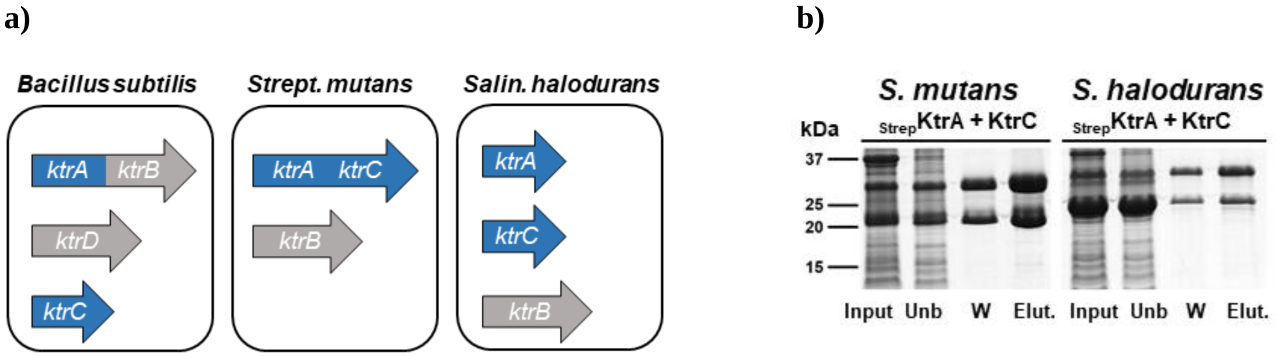


**Figure S4: Heteromeric assembly of RCK protein from other bacterial species. a)** Cartoon representations of Ktr gene organizations in *Bacillus subtilis*, *Streptococcus mutans* and *Salinococcus halodurans*. **b)** SDS-PAGE analysis of Streptactin pull-down experiments from lysates of *E. coli* co-expressing pairs of RCK proteins encoded in the genomes of *Streptoccocus mutans* and *Salinococcus halodurans*. One of the RCK proteins is tagged with a Strep tag. Cell lysate (input), unbound (unb), wash (W) and eluted (Elut.) fractions.

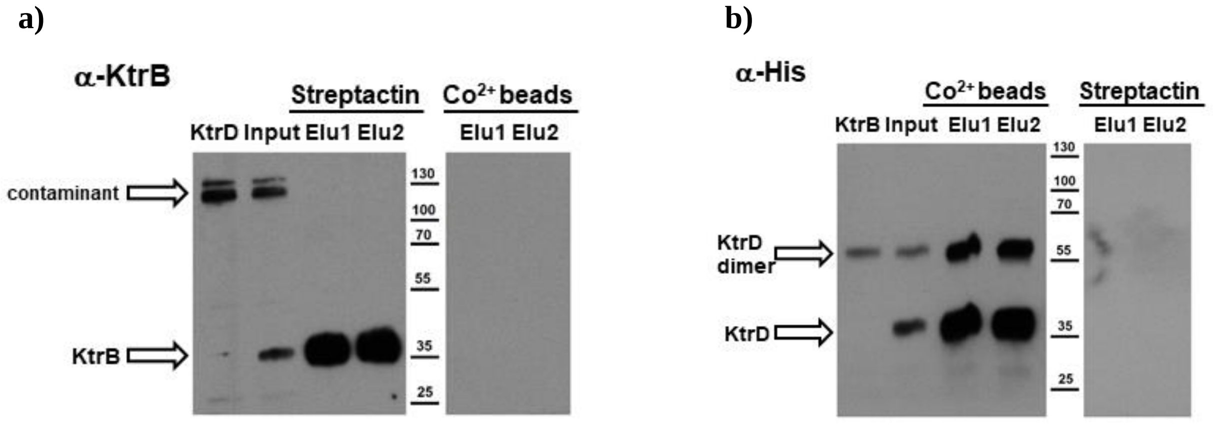


**Figure S5: KtrB and KtrD do not form heteromeric membrane protein dimers.** Detergent extracted lysates of *E. coli* cells co-expressing Strep tagged KtrB ($_{Strep}$KtrB) and His tagged KtrD ($_{His}$KtrD) were submitted to pulldown experiments with **a)** Streptactin beads or **b)** $Co^{2+}$ beads. Eluted proteins were analyzed by western blots with antibodies against **a)** KtrB (α-KtrB) or **b)** His tag (α-His), revealing the presence of $_{Strep}$KtrB in the elution from Streptactin beads but not in $Co^{2+}$ beads and of $_{His}$KtrD in elution from $Co^{2+}$ beads but not in Streptactin beads. KtrB and KtrD have similar migration pattern in SDS-PAGE and are sometimes observed as monomers and dimers in SDS-PAGE. Purified proteins are included as controls.

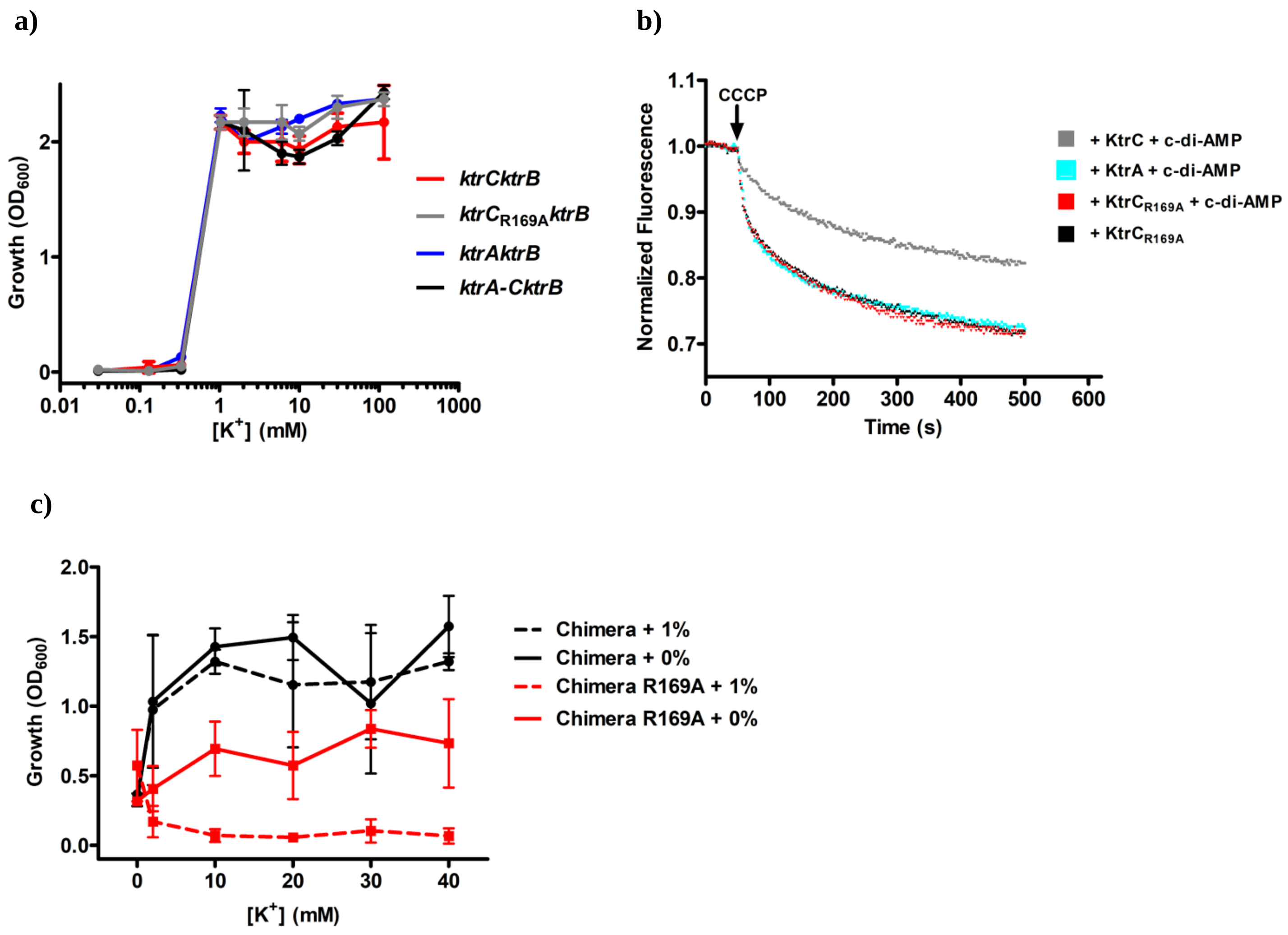


**Figure S6: Properties of RCK proteins with altered c-di-AMP binding sites. a)** Overnight growth, measured by optical density at 600 nm, of TK2420 *E. coli* strain co-expressing KtrB with KtrA, $KtrC_{R169A}$ or KtrA/KtrC chimera (KtrA-C) in different $K^+$ concentrations. **c)** Pyranine fluorescence flux curves for Ktr channels assembled by addition of KtrA, KtrC or $KtrC_{R169A}$ to KtrB-reconstituted liposomes in the presence of ATP and $Mg^{2+}$ with and without 20 µM c-di-AMP. **d)** $K^+$ requirement assays for JH642 Δ*ktrA*Δ*ktrC* strain transformed with pTH1xp plasmid encoding the KtrA/KtrC chimera or the KtrA/KtrC chimera with equivalent mutation to R169A in KtrC with (1% xylose) and without induction (0% xylose) of protein expression. Mean ±SD for n=3.

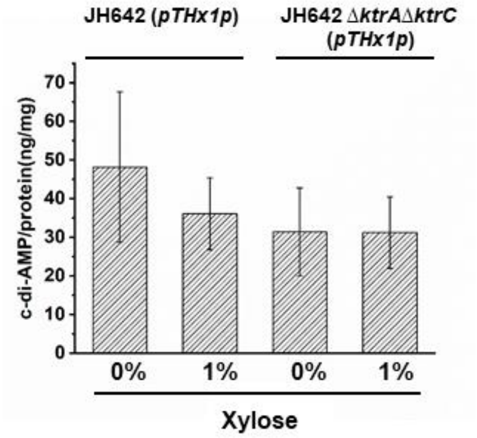


**Figure S7: c-di-AMP levels in *B. subtilis* JH642 strain.** Cultures of JH642 and JH642 Δ*ktrA*Δ*ktrC* strains transformed with the empty pTHx1p plasmid were exposed to 0% or 1% xylose and their c-di-AMP levels were determined, showing no effect of xylose exposure. Mean ±SD for n=3.

**Table S1: Strains and plasmids**

| Strains/plasmids | Relevant characteristics | References |
|---|---|---|
| ***E. coli* strains** | | |
| XL1-Blue | *recA1 endA1 gyrA96 thi-1 hsdR17 supE44 relA1 lac* [F´ *proAB lac*Iq*Z*Δ*M15* Tn*10* (Tetr) | Laboratory collection |
| ***B. subtilis* strains** | | |
| 168 | *trpC2* | Laboratory collection |
| JH642 | *pheA1*; *trpC2*; *citS642* | Laboratory collection |
| Δ*ktrA* | Δ*ktrA* deletion mutant of *B. subtilis* strain JH642 | This study |
| Δ*ktrB* | Δ*ktrB* deletion mutant of *B. subtilis* strain JH642 | This study |
| Δ*ktrC* | Δ*ktrC* deletion mutant of *B. subtilis* strain JH642 | This study |
| Δ*ktrD* | Δ*ktrD* deletion mutant of *B. subtilis* strain JH642 | This study |
| Δ*ktrD*Δ*ktrB* | Δ*ktrD* Δ*ktrB* deletion mutant of *B. subtilis* strain JH642 | This study |
| Δ*ktrA*Δ*ktrC* | Δ*ktrA* Δ*ktrC* deletion mutant of *B. subtilis* strain JH642 | This study |
| Δ*kimA*Δ*ktrA* | Δ*kimA* Δ*ktrA* deletion mutant of *B. subtilis* strain *168* | This study |
| Δ*kimA*Δ*ktrC* | Δ*kimA* Δ*ktrC* deletion mutant of *B. subtilis* strain *168* | This study |
| Δ*ktrB*-AB | Δ*ktrB* strain with *ktrAB* operon integrated in *amyE* | This study |
| Δ*ktrA*Δ*ktrC*-C | Δ*ktrA*Δ*ktrC* strain with *ktrC** operon integrated in *amyE* | This study |
| Δ*ktrA*Δ*ktrC*-C$^{R169A}$ | Δ*ktrA*Δ*ktrC* strain with *ktrC**$^{R169A}$ operon integrated in *amyE* | This study |
| Δ*ktrA*Δ*ktrC*-A | Δ*ktrA*Δ*ktrC* strain with *ktrAB*$^{T59stop/A64stop}$ operon integrated in *amyE* | This study |
| JH642(pTH1xp) | JH642 strain with the replicative plasmid pTH1xp | This study |
| Δ*ktrA*Δ*ktrC*(pTH1xp) | Δ*ktrA*Δ*ktrC* strain with replicative plasmid pTH1xp | This study |
| Δ*ktrA*Δ*ktrC*(pTH1xp-*ktrC*) | Δ*ktrA*Δ*ktrC* strain with replicating plasmid pTH1xp-*ktrC* | This study |
| Δ*ktrA*Δ*ktrC*(pTH1xp-*ktrC*$^{R169A}$) | Δ*ktrA*Δ*ktrC* strain with replicative plasmid pTH1xp-*ktrC* $^{R169A}$ | This study |
| Δ*ktrA*Δ*ktrC*(pTH1xp-chimera) | Δ*ktrA*Δ*ktrC* strain with replicative plasmid pTH1xp-chimera | This study |
| Δ*ktrA*Δ*ktrC*(pTH1xp-chimera$^{R169A}$) | Δ*ktrA*Δ*ktrC* strain with replicative plasmid pTH1xp-chimera$^{R169A}$ | This study |
| **Plasmids** | | |
| pDG364 | Cm$^R$, plasmid for integration at *amyE* locus in *B. subtilis* chromosome | BGSC |
| pDG1730 | Spec$^R$, plasmid for integration at *amyE* locus in *B. subtilis* chromosome | BGSC |
| pDG364-*ktrAB*_1 | pDG364 derivate plasmid for integration of 3´region of *ktrAB* operon | This study |

| | | |
|---|---|---|
| pDG1730-*ktrAB*_2 | pDG1730 derivate plasmid for integration of 5´region of *ktrAB* operon | This study |
| pDG364-*ktrAB*$^{T59stop/A64stop}$ | pDG364-*ktrAB*_1 derivative plasmid with 2 stops codons at codons for amino acids T59 and A64 in KtrB | This study |
| pDG364-*ktrC** | pDG364-*ktrC* derivative plasmid with a single amino acid exchange in KinC (H224A) and four stop codons at codons for amino acids L5, F6, Y130 and F131 of YkqA | This study |
| pDG364-*ktrC**$^{R169A}$ | pDG364-*ktrC* derivative plasmid with a single amino acid exchange in KinC (H224A), four stop codons at codons for amino acids L5, F6, Y130 and F131 of YkqA and R169A exchange in KtrC | This study |
| pTH1xp | $Cm^R$, *E. coli*/*B.* spp. shuttle vector for xylose inducible expression | (4) |
| pTH1xp-*ktrC* | pTH1xp derivative plasmid for overexpression of KtrC | This study |
| pTH1xp-*KtrC*$^{R169A}$ | pTH1xp-*ktrC* derivative plasmid for overexpression of KtrC with mutation R169A | This study |
| pTH1xp-*ktrA* (ATG) | pTH1xp derivative plasmid for overexpression of KtrA. The start codon was changed from TTG to ATG | This study |
| pTH1xp-chimera | pTH1xp derivative plasmid for overexpression of RCK chimera fusing the KtrA N-terminal region (amino acids 1-125) and the KtrC C-terminal region (amino acids 122-221) | This study |
| pTH1xp-chimera$^{R169A}$ | pTH1xp derivative plasmid for overexpression of RCK chimera fusing the KtrA N-terminal region (amino acids 1-125) and the KtrC C-terminal region (amino acids 122-221) with mutation R169A in KtrC | This study |

$Cm^R$ – chloramphenicol resistance; $Spec^R$ – spectinomycin resistance


1. J. Gundlach *et al.*, Sustained sensing in potassium homeostasis: Cyclic di-AMP controls potassium uptake by KimA at the levels of expression and activity. *J Biol Chem* **294**, 9605-9614 (2019).
2. Y. Oppenheimer-Shaanan, E. Wexselblatt, J. Katzhendler, E. Yavin, S. Ben-Yehuda, c-di-AMP reports DNA integrity during sporulation in Bacillus subtilis. *EMBO Rep* **12**, 594-601 (2011).
3. B. M. Wendel *et al.*, A Central Role for Magnesium Homeostasis during Adaptation to Osmotic Stress. *mBio* **13**, e0009222 (2021).
4. M. Irla *et al.*, Genome-Based Genetic Tool Development for Bacillus methanolicus: Theta- and Rolling Circle-Replicating Plasmids for Inducible Gene Expression and Application to Methanol-Based Cadaverine Production. *Front Microbiol* **7**, 1481 (2016).
